# Impaired ovarian development in a zebrafish *fmr1* knockout model

**DOI:** 10.1101/2024.02.10.579749

**Authors:** Rita Rani, N Sushma Sri, Raghavender Medishetti, Kiranam Chatti, Aarti Sevilimedu

## Abstract

Fragile X syndrome (FXS) is an inherited neurodevelopmental disorder and the leading genetic cause of autism spectrum disorders. FXS is caused by loss of function mutations in Fragile X mental retardation protein (FMRP), an RNA binding protein that is known to regulate translation of its target mRNAs, predominantly in the brain and gonads. The molecular mechanisms connecting FMRP function to neurodevelopmental phenotypes are well understood. However, neither the full extent of reproductive phenotypes, nor the underlying molecular mechanisms have been as yet determined. Here, we developed new *fmr1* knockout zebrafish lines and show that they mimic key aspects of FXS neuronal phenotypes across both larval and adult stages. Results from the *fmr1* knockout females also showed that altered gene expression in the brain, via the neuroendocrine pathway contribute to distinct abnormal phenotypes during ovarian development and oocyte maturation. We identified at least three mechanisms underpinning these defects, including altered neuroendocrine signaling in sexually mature females resulting in accelerated ovarian development, altered expression of germ cell and meiosis promoting genes at various stages during oocyte maturation, and finally a strong mitochondrial impairment in late stage oocytes from knockout females. Our findings have implications beyond FXS in the study of reproductive function and female infertility. Dissection of the translation control pathways during ovarian development using models like the knockout lines reported here may reveal novel approaches and targets for fertility treatments.

Graphical Abstract

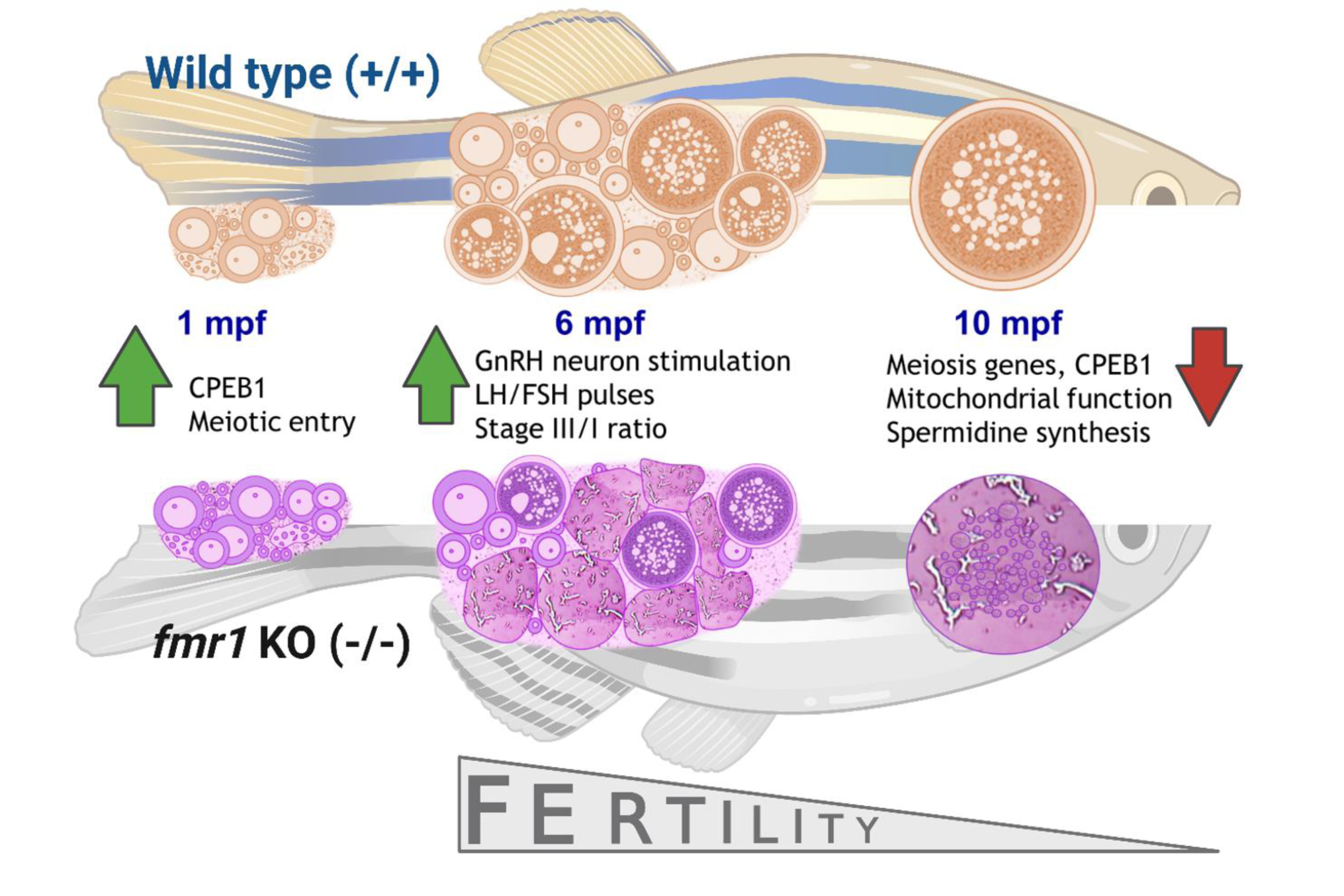

## Introduction

Fragile X syndrome (FXS, OMIM 300624) is a heritable intellectual disability disorder that is caused by the absence of the Fragile X mental retardation protein (FMRP) (Salcedo-Arellano et al., 2020). Patients with FXS display varying degrees of intellectual deficits, macroorchidism and distinct facial dysmorphisms including an elongated face and long ears. The disease mechanism of FXS has been studied extensively, both in pre-clinical models as well as clinical setting, as it is a dominant monogenic cause of autism. FMRP is expressed in all neuronal cell types, where it binds numerous transcripts implicated in autism, including those involved in cell signaling, synaptic development and function, axonal and dendritic development, and serves as a translation modulator (Richter and Zhao, 2021). In FXS patients, a loss of stimulus dependent translation regulation of these key mRNAs is thought to be the molecular basis of the neuronal phenotypes. However, therapies aimed at these molecular mechanisms were not found to be efficacious in clinical trials (Santoro et al., 2012). This may be explained partly based on the time of intervention, as FMRP is known to play a role in early neuronal development. It is further supported by a proof-of-concept evidence that neurodevelopmental disorders, such as FXS, may be responsive to early pharmacological intervention (Asiminas et al., 2019; Medishetti et al., 2019).

FMRP is highly expressed in the gonads, the gametes and ubiquitously in the embryo during early development (Malecki et al., 2020; Takahashi et al., 2015) where it likely plays a key role in orchestrating the temporally and spatially controlled translation of maternal mRNAs, along with other RNA binding proteins (RBPs) (Rosario et al., 2017). Therefore, it is expected that a loss of this key regulator is likely to significantly impact fertility, oocyte maturation and/or early development of the embryo. This is important to study because it is probable that at least some of the neurodevelopmental defects and behavioral phenotypes seen in adults originate from this early disruption of the translation program (Krisher, 2004). Also important to note that primary ovarian insufficiency is a well reported Fragile-X associated condition, however it is associated with pre-mutation at the FMRP locus, and likely caused by *fmr1* RNA toxicity, and not reported to be caused by a reduction or loss of FMRP (Man et al., 2017)

Several animal models have been generated to understand the function of FMRP, including knockouts in the fruitfly, zebrafish, mouse and rat. However, there are only a few studies that focus on its role in oogenesis and early embryonic development: several in Drosophila, which reveal the role of *dfmr1* in early oocyte development (Costa et al., 2005; Deshpande et al., 2006; Epstein et al., 2009; Monzo et al., 2010; Yang et al., 2007), and a few in mice that suggest a role in fertility (Mok-Lin et al., 2018; Villa et al., 2023). However, a longitudinal study investigating FMRP function at different stages of ovarian development and possible mechanisms of the observed phenotypes, has not yet been reported in a vertebrate model. The previously reported zebrafish *fmr1* knockout lines mimic the neuronal phenotypes seen in FXS patients, reinforcing the idea that zebrafish is an excellent vertebrate model to study FMRP function. Therefore, in our study, we used zebrafish *fmr1* knockout lines to investigate the impact of FMRP loss on reproduction and fertility at various ages. We found a delayed reproductive dysfunction phenotype in the knockout females and identified a few different molecular mechanisms contributing to this specific phenotype.

## Materials and Methods

### Animal Ethics Procedures and Approval

All experiments with zebrafish were done in a CCSEA-approved zebrafish facility at Dr. Reddy’s Institute of Life Sciences (1100/po/Re/s/07/CPCSEA) in Hyderabad, India. The facility also has US-NIH OLAW assurance (F22-00539). All procedures and protocols were reviewed and approved by the Institutional Animal Ethics Committee (Protocol approval DRILS/IAEC/AS/2019-1). The “Guidelines for Experimentation on Fishes, 2021” published by CPCSEA was used as a reference.

### Zebrafish culture, maintenance and breeding

The origin of the zebrafish used was a group of Indian wild-caught fish obtained from a commercial breeder. These fish were grown and bred in-house over more than 3 years. The fish were maintained in a semi-automated housing system (ZebTEC rack with active blue technology, Tecniplast, Italy). This system circulates filtered and aerated water to all fish tanks and maintains a stable pH of 7.0, a temperature of 28°C, and conductivity of 500μS. A photoperiod of 14h light and 10 h dark was followed. The fish were fed three times every day with either dry or live feed (pellets or live brine shrimp hatched in-house) with live feed at least once daily. Adult fish between three months and twelve months of age were bred for experiments. Breeding tanks (1.7L) were used to set up pairs of male and female fish on the evening prior to the breeding day. Zebrafish breeding is responsive to photoperiodic cues and they lay eggs or sperm at first light. Typical breeding pairs in our facility produce an average of 300 eggs with an 80% fertilization rate. Unfertilized or non-viable eggs were discarded and fertilized eggs were transferred into a fresh Petri dish, washed at least two times in system water and maintained in a 28°C incubator.

### sgRNA synthesis

Target regions for CRISPR mediated editing for *fmr1* were chosen using Benchling software in exons 3 and 4. Single guide RNAs were synthesized in-house using standard protocols as described previously (Wierson et al., 2020). Briefly, sgRNA was ordered as single stranded DNA with the addition of T7 promoter at 5’end and tail oilgo sequence at 3’end (Table S1). DNA template for sgRNA synthesis was generated by standard overlap PCR with the sgRNA and tail oligos, and used for in-vitro transcription using the HiScribe™ T7 High Yield RNA Synthesis Kit (NEB). Finally, sgRNA was treated with DNase (NEB) and purified using the RNA Clean and Concentrator kit (Zymo Research) according to manufacturer’s instructions. sgRNA concentration was measured using a Biospectrophotometer (Eppendorf) and integrity was checked on the urea PAGE.

### Microinjection and generation of FMR knockout lines

CRISPR/Cas9 mediated knockout generation was performed as described previously (Medishetti et al., 2022). Briefly, one cell stage zebrafish embryos were injected with 5nl of Cas9-sgRNA mix (400ng of Cas9 protein and 0.2-0.5µg of sgRNA in 5µl of 10mM Tris pH 8.0). After 24hpf, dead and abnormal embryos were removed and remaining were raised to adulthood as F0 founders. Sexually mature F0 founders were outcrossed and F1 offspring were screened for indels by tail-clip PCR using the standard heteroduplex mobility assay (HMA) with genotyping primers (Table S1). Positive F1 offspring were raised to adulthood, F1s of the same type were interbred to generate F2, and homozygotes were sequenced to identify the edited allele. For genotyping, 24hpf embryos or tail fin clip was heated at 95^0^C for 30 mins in 45 µL of 50mM NaOH solution and followed by neutralization with 5µL of 1M Tris pH 8.0. After a short spin, 0.3ul was used as input for a 5ul PCR reaction. PCR reaction was set up with 2X Q5 mix and genotyping primers. PCR product was run on 10% NATIVE PAGE.

### Locomotion and behaviour analysis

Locomotion and behaviour analysis in larvae was performed as reported previously for manual recording (Medishetti et al., 2019). For automated behaviour analysis, the Zebrabox system and Zebralab software were used. Single larvae (7, 10, 14dpf) were gently placed in a 24 well plate (in 1ml of E3) 1hr before the experiment and placed inside the Zebrabox for acclimatization for 10 minutes. For the light dark locomotion assay, locomotion was recorded over 10min dark, followed by alternative light/dark cycles of 3 minutes each. The integration period was set to 1 min; total distance was calculated for each minute. The data obtained was analysed and graphed using Microsoft Excel, and reported as reported as mean ± SEM (n=60 (7dpf), n=15 (10dpf), n=90 (14dpf)).

Behaviour assays in adult male zebrafish were performed manually, as reported in (Kalueff, 2017) with minor modifications described below, and analysed using AnyMaze software. The light dark test was conducted in an aquarium of size 220 × 210× 140 mm, separated into two chambers, one of which was covered with the white paper and the other with matte black paper covering it on all sides of the tank. A camera pointed towards the light chamber with the LED light panel was used to record for 10 minutes at 30 frames per second covering the whole area of the tank. The tank was filled with 5 litres of water that included 1.2 g of sea salt for conductivity. The fish were acclimatized in the behaviour room for 1hr, and then introduced into light dark box and recorded for 10min. The percentage time spent in the dark zone was measured. The novel tank was conducted in a tank apparatus consisting of a narrow tank divided horizontally into a top and bottom zone. A camera pointed vertically towards the chamber was used to record for 10 minutes at 30 frames per second. The tank was filled with 5 litres of water that included 1.2 g of sea salt for conductivity. The fish were acclimatized in the behaviour room for 1hr, and then introduced into the test tank and recorded for 10min. The percentage time spent in the bottom zone was measured. The open field test (OFT) was conducted in the OFT Box (220 ×210 ×140 mm) with a defined inner zone and outer zone. The fish were acclimatized in the behaviour room for 1hr, and then introduced into the test tank and recorded for 10min. The percentage time spent in the outer zone was measured. The data from these experiments were analysed using Graphpad PRISM and reported as mean +/-SEM (statistical test One-way ANOVA).

### Gene expression analysis: RNA extraction and quantitative real-time PCR

RNA was isolated from different tissue (7dpf larvae, adult brain, oocytes and ovary) using Trizol (NEB) and purified using the RNA Clean and Concentrator kit (Zymo Research) according to manufacturer’s instructions. For tissue (brain and ovary) homogenization was done with RNase free plastic pestle and larvae were homogenized using a 27G syringe. cDNA was made using 500ng RNA with PrimeScript RT Reagent Kit (TaKaRa). Real time PCR was set up using TB Green Premix Ex Taq II (Tli RNase H Plus) (TakaRa) in a Quantstudio5 machine. RNAPD (*polr2d*) was used as the reference gene. Sequences of all the primers used in the study are listed in Table S1. The changes in RNA expression were quantified by the ΔΔCT method, normalized to the internal control, and then, in certain experiments, to the control group for each experiment. The results from three or more independent experiments were averaged and are reported, with error bars representing S.E.M. Statistical analysis (t-test) was performed using Graphpad PRISM software.

### Protein measurement by Western Blot

Zebrafish larvae or adult brain were lysed in RIPA lysis buffer with protease inhibitor cocktail (10 μg/mL each aprotinin and leupeptin). Protein concentration was determined by BCA method. Approximately 30µg protein was loaded on 10% SDS-PAGE and then transferred to PVDF membrane (Millipore) followed by blocking, primary (1:1000-5000) and secondary antibody (1:5000-10,000) incubation. Antibodies used are listed in Table S2. To probe for multiple proteins, the membrane was cut at the appropriate size marker. Signals were detected with ECL western blotting substrate (TaKaRa) and image was acquired using Azure Biosystems. Images were quantified by using Image J. Alpha-tubulin was used as the reference gene for protein measurements.

### Immunohistochemistry

IHC was performed as described in (Hammond-Weinberger and Zeruth, 2019). Briefly, stage III oocytes (∼0.4-0.7mm diameter, (Selman et al., 1993)) were collected from adult zebrafish (n=3/group) after dissection, fixed in 4% PFA overnight and then washed for 10mins (5 times) with PBST. Oocytes were dehydrated and then rehydrated with methanol followed by blocking (10% horse serum in PBST) for 4hrs at room temperature. They were incubated with primary antibody (1:200 in blocking solution) for 3 days at 4^0^C and then washed 5 times (20mins) with PBST at room temperature and incubated with the appropriate Alex Flour-488 conjugated secondary antibody for 2 days at 4^0^C. Oocytes were washed 5 time (20mins) with PBST at room temperature and then mounted in mounting media. Images of 5-10 oocytes/n were taken using EVOS cell imaging system.

### PCR polyadenylation tail (PAT) assay

PCR polyadenylation tail (PAT) assay was performed as described previously (Sallés and Strickland, 1995). Briefly, 250ng RNA after DNase treatment was heat denatured in the presence of 56ng of phosphorylated oligo (dT) in 7µl volume at 65^0^C for 5mins and immediately placed at 42^0^C. 13µl of pre warmed master mix (4µl of 5 X RT buffer, 2µl of 0.1M DTT, 2µl of 2.5mM dNTPS, 0.1µl of 100mM ATP and 2.5µl of T4 DNA ligase (40U/µl) and 2.4µl of H_2_0) was added and incubated at 42^0^C for 30mins. Then 560ng of anchor oilgo(dT) was added and reaction was incubated at 12^0^C for 2hrs. After 2 hrs 1µl of Reverse Transcriptase (PrimeScript RT, Takara) was added and reverse transcription was performed at 42^0^C for 1hr followed by inactivation at 70^0^C for 30 mins. PCR was performed using a gene specific forward primer (200-600bp upstream of the poly(A) addition site) and anchor T reverse primers using standard PCR programme. PCR products were visualized by native PAGE (12% gel). PAT-PCR was performed in oocytes collected from n=3 fish/group. Representative gel images are shown.

### H&E staining and Histology

Female zebrafish (n=3/group) were collected three days after the last breeding event and sampled for histopathological examination. Briefly, the fish were fixed in Davidson fixative solution and dehydrated through increasing ethanol series. Then the fish were embedded after immersion in liquid paraffin at 60 °C for 2h. The Formalin-Fixed Paraffin Embedded (FFPE) were sectioned at 3 μm-thickness, and then rehydrated using Xylene and Ethanol solutions. Finally, fish sections were stained with Hematoxylin and Eosin for morphological analysis. Slides were observed using a Zeiss Primovert microscope at 4X, and images were acquired using Axiocam 208 (colour).

## Results

### Generation of the fmr1 knockout (KO) zebrafish lines

We used the standard CRISPR workflow to generate *fmr1* knockout zebrafish lines (Medishetti et al., 2022). Single guide RNAs were designed to target early exons (3 and 4) of zebrafish *fmr1*, in order to maximise the likelihood of complete loss of function (Fig. 1A). In an *in vitro* catalytic activity assay, both guides were able to direct Cas9 mediated dsDNA cleavage of the target DNA amplicon (Fig. S1A). Cas9-RNP complexes were assembled with both guides and used for editing the *fmr1* gene *in vivo*. Subsequently, standard protocols were followed to identify two edited alleles (Fig. S1B), generate heterozygotes, and homozygotes. Two independent lines were maintained, each carrying a different mutant allele of *fmr1*: type 6 contained a 13bp insertion and type 11 had a 4bp deletion (5bp deletion and 1bp insertion), each expected to result in a premature termination of translation in exon 4 (Fig. 1B, S1C). The levels of *fmr1* mRNA were found to be significantly reduced (Fig. S2A), and protein levels were undetectable (Fig. S2B) in the knockout zebrafish. Both lines are henceforth referred to as *fmr1* knockout (KO) zebrafish lines.

**Figure 1:**
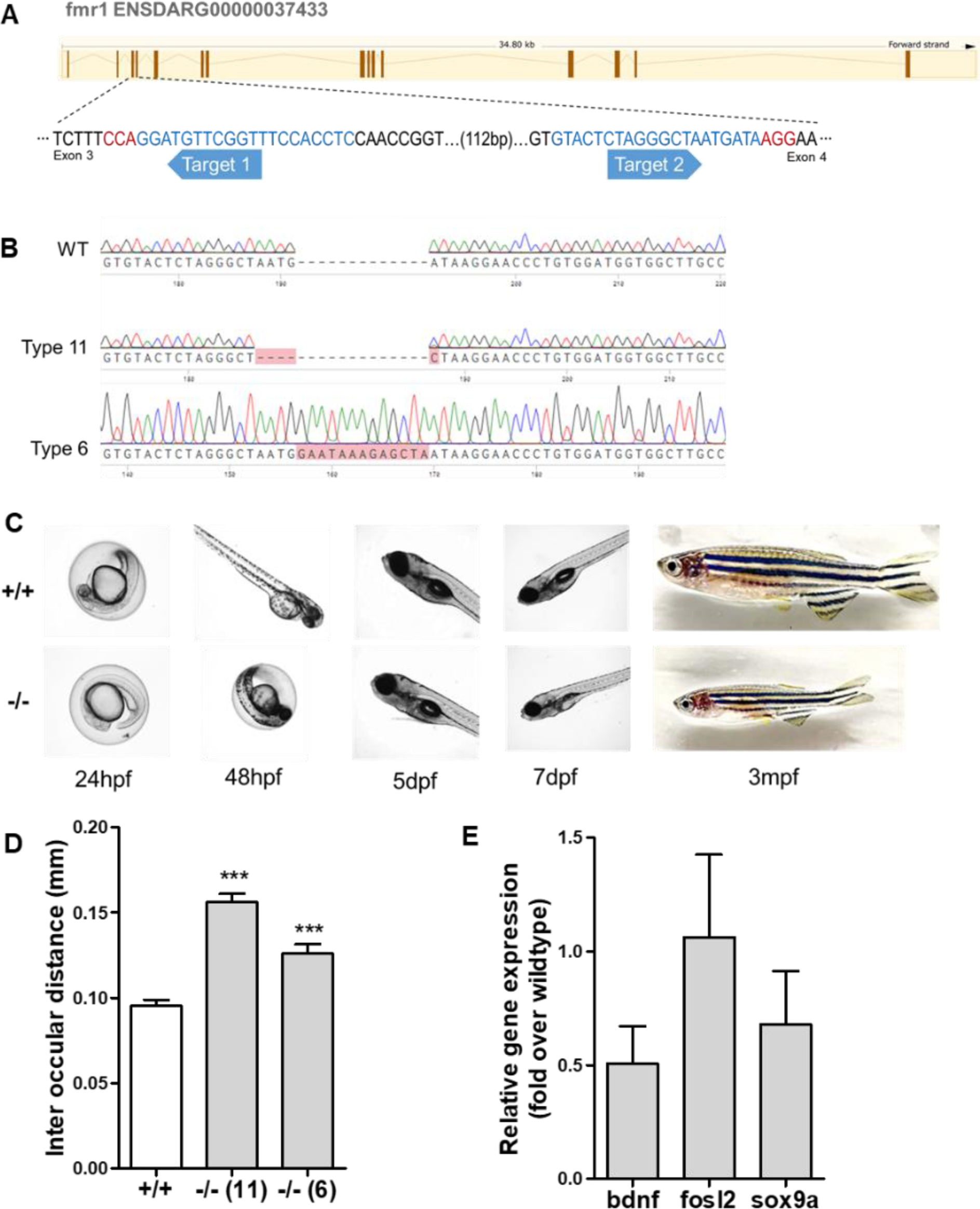
Characterization of the *fmr1* KO zebrafish line: A. Location of single guide RNAs at the *fmr1* locus. B. Sequence of the edited alleles in Type 6 and Type 11 KO lines. C. Imaging of the KO larvae at various time points during development D. Quantification of the interocular distance, from Alcian blue stained larvae, showing increased spacing in the KO larvae (n=6). E. Relative expression levels of genes involved in craniofacial development showing a slight decrease in the KO larvae at 7dpf (n=3).

Heterozygote crosses involving *fmr1* mutant alleles resulted in progeny, which followed Mendelian ratios at birth. Even though their development appeared to be similar to wild type, the rate of survival of knockout zebrafish to adulthood (4mpf) was less than expected (16 +/-6% S.D, as compared to 25%), indicating lower fitness of the complete knockout. The *fmr1* knockout zebrafish were fertile but more erratic in breeding than the age matched wildtype zebrafish. Early development of knockout embryos appeared to be marginally slower than wildtype for the first 24-48hpf (Fig. 1C). In the first two weeks, the knockout larvae were smaller and ∼10-20% had physical malformations, such as, lack of air sac and craniofacial abnormalities (jaw defects). By 3mpf, most knockout fish looked similar to wildtype, but about 20% were smaller and devoid of operculum (Fig 1C). Minor changes in the craniofacial architecture were observed in the knockout larvae by Alcian blue staining (Fig. S3), similar to previous reports (Hu et al., 2020; Medishetti et al., 2019; Tucker et al., 2006), the most prominent being an increase in inter-ocular space (Fig. 1D).

### Behavioural phenotypes in the fmr1 knockout larvae

Patients with FXS display one or more of a wide range of behavioural symptoms such as increased anxiety, hyperactivity, cognitive deficits, irritability, and sleep disruptions (Clinical Synopsis - #300624 - FRAGILE X SYNDROME; FXS - OMIM). We had previously demonstrated evidence of increased anxiety, reduced fear cognition and increased irritability in the transient *fmr1* mRNA knockdown larval model (Medishetti et al., 2019). However, in the *fmr1* KO larvae, we did not detect a significant increase in anxiety or irritability, or decrease in fear cognition, measured by the same assays. In order to detect subtle phenotypes, Viewpoint Zebrabox and Zebralab system was used to perform automated locomotion tracking in a much large number of knockout larvae (48-96 larvae per group) at various stages during their growth. We found a robust hyperactivity phenotype at 14dpf, which was not apparent at 7dpf or 10dpf (Fig. 2A, Fig.S4 A, B). However, we did not measure a robust and significant increase in the thigmotaxis behaviour (a measure of anxiety driven by a novel environment) in the larval assays even at 14dpf. However, in the KO larvae we observed a much higher rate of locomotion response in response to a light-to-dark or dark-to-light transition (Fig. S4C, D), which is supportive of an increase in stimulus driven anxiety in these larvae.

**Figure 2:**
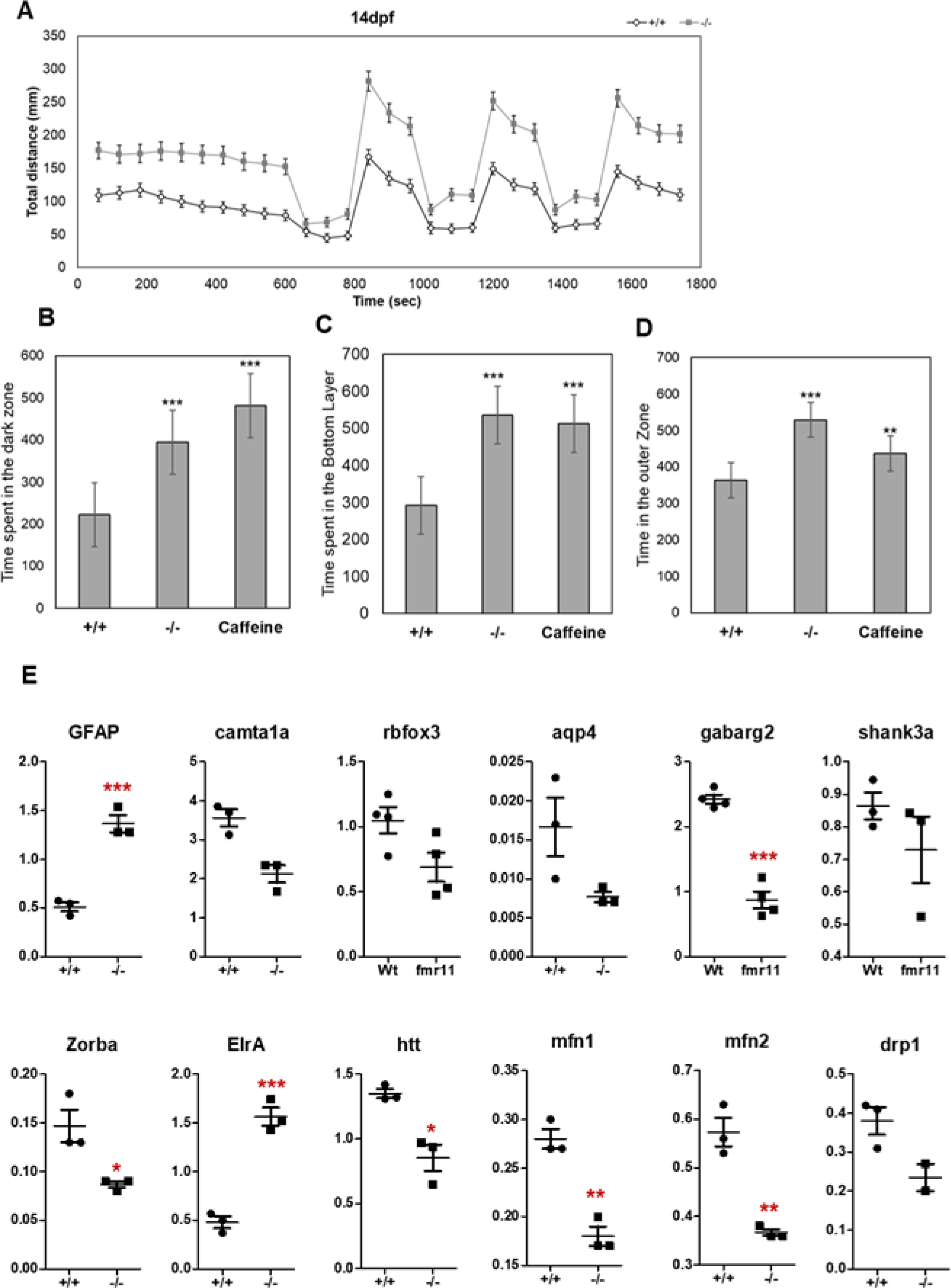
Behavioural phenotypes and molecular changes in the KO larvae and adult zebrafish: A. Locomotion tracking in individual larvae showing hyperactivity in the KO larvae at 14dpf (n=90/group). B-D Behavioral assays in adult zebrafish (6-9mpf) B. Light dark preference test: Percentage time spent in dark zone C. Novel tank test. Percentage time spent in bottom zone. D. Open field test: Percentage time spent in outer zone. Caffeine treated fish were used as a positive control for increased anxiety (n=6/group). E. Relative expression levels of various groups of genes as described in the text (n=4/group).

Since we found a difference in the behavioural phenotypes exhibited by the KO larvae (ours as well as previously published reports (den Broeder et al., 2009; Hu et al., 2020)), and the larvae with a transient mRNA knockdown (Medishetti et al., 2019; Tucker et al., 2004), we tested if this was due to a compensation from the upregulation of other *fmr1* like genes *fxr1* and *fxr2.* While we observed a robust decrease in *fmr1* mRNA levels at 7dpf in the KO larvae, we did not observe a significant increase in the expression of either *fxr1* or *fxr2* (Fig S2A). And, as previously reported, we observed a decrease in the levels of *bdnf* (neuronal development), and moderate but reproducible decreases in *fosl2*, and *sox9a* (contributing to the mild craniofacial deformities) (Fig. 1E). Among all of the other markers of early neuronal development, the only other significant changes observed were a robust increase in GFAP, a marker of astrocytes and a decrease in olig2, the oligodendrocyte marker (Fig S5A).

A major theory in the FXS field postulates that mGluR5 upregulation is the key basis of the pathomechanism of FXS (Bear et al., 2004). In the absence of FMRP, which regulates the translation of mGluR5, the mGluR5 protein level and the downstream signalling cascade is upregulated, leading to stimulus independent signalling and excitotoxicity. Therefore, to test this hypothesis, we measured the levels of mGluR5 and other members of the signalling cascade (p-ERK and p-Akt) in the KO and matched wildtype larvae, and saw no significant increase in these markers (Fig. S5B). This is in contrast to what we observed when *fmr1* mRNA was transiently knocked down (Medishetti et al., 2019), where mGluR5 as well as the downstream signalling was upregulated. In the case of the knockout, it is possible that there may be a compensatory pathway of mGluR5 translation regulation in zebrafish that is yet to be characterized, or localized changes in expression or activation, which were not captured in these experiments.

### Behavioural phenotypes and altered gene expression in the fmr1 knockout adult zebrafish

The behaviour phenotypes and brain-specific gene expression were assessed in the adult KO zebrafish (>6-12mpf, males). Adult behaviour was recorded and analysed using AnyMaze software. Three established paradigms were used (Kalueff et al., 2013). The light-dark preference test, which relies on the innate preference of adult zebrafish for the dark environment. Any increase in anxiety results in the fish spending more time in the dark side. The “novel tank diving” (NTD) test which relies on the zebrafish’s innate propensity to dive, freeze, and limit exploration in novel settings. Increased anxiety results in increased amount of time spent in the deeper zones. The open field test (OFT) to measure thigmotaxis, which is a typical method for evaluating exploratory behaviour and general activity in zebrafish. Increased anxiety or intellectual deficits will result in reduced exploration.

In each of these tests, the KO fish spent more time in the dark zone (Fig. 2B), the bottom layer (Fig. 2C) and the outer zone (Fig. 2D), much like caffeine treated wild-type fish, which demonstrates an increased level of anxiety and a reduced tendency to perform exploratory activity in the KO fish (an indirect indicator of intellectual abilities). However, very similar to what was observed in the KO larvae, we saw no upregulation in the mGluR5 protein levels, or activity as measured by p-ERK and p-Akt levels in the brain lysates made from adult KO zebrafish (Fig. S6A, B and C). It is possible that these changes occur in specific regions or cell types in the brain, and are therefore not measurable in an assay performed on the whole brain lysate. These results suggest that the observed phenotypes may be mediated by changes that occur independent of the mGluR5 pathway.

We carried out gene expression analysis in the brain tissue of KO and age matched wildtype fish, for three classes of genes a) FMRP targets (mRNAs directly bound by FMRP or mRNAs whose polyadenylation profile/translation is altered in the absence of FMRP) b) genes associated with autism c) genes whose expression has been reported to be changed in other FXS models or patients (Table S3). We found significant changes in neuronal markers indicative of increased astrocytosis (GFAP), reduced neuronal number and cognition (*rbfox3* and *camta1a*), BBB integrity (*aqp4*) and significant decrease in the inhibitory neurotransmitter receptor GABA-A(2) (*gabarg2*). We also found a reduction in neuronal translation control factors (CPEB1(*zorba*) and the ELAVL family protein, *elrA*). Consistent with recent reports suggesting that Huntingtin (*htt*) is a direct mRNA target of FMRP (Shen et al., 2019), we found a decrease in *htt* levels in the KO zebrafish brain, and a corresponding change in several genes involved in mitochondrial dynamics indicative of reduced fusion (Fig 2E). A decrease in protein levels of Mfn2, Opa1 and Zorba was also detected in the KO brain tissue (Fig S6D-F). This mitochondrial dysfunction and persistent astrocyte activation in the FXS brain, could together be the basis for impaired neuronal development and function manifested in the form of altered behavioural responses in these KO zebrafish.

In contrast to the drastic decrease in the expression of the GABA-A receptor *gabarg2* observed in the KO male brain (old fish (Fig.2E) and young fish (Fig 3A)), the expression levels of this receptor were unchanged in the young female KO brains (Fig 3B). In the sexually mature adult brain, GABA acts as an excitatory neurotransmitter for GnRH neurons (Seong-Kyu et al., 2002; Watanabe et al., 2014) eliciting the release of GnRH, which subsequently promotes release of gonadotropins (LH and FSH) from the pituitary. In addition, in the brains of sexually mature female KO fish, we detected a reproducible increase in the expression of the neuroendocrine hormones *gnrh3,* and *fsh* (Fig 3C). These findings support the idea of altered hypothalamic-pituitary control of reproductive development in the female KO fish, similar to that reported in the female KO mice (Villa et al., 2023).

**Figure 3:**
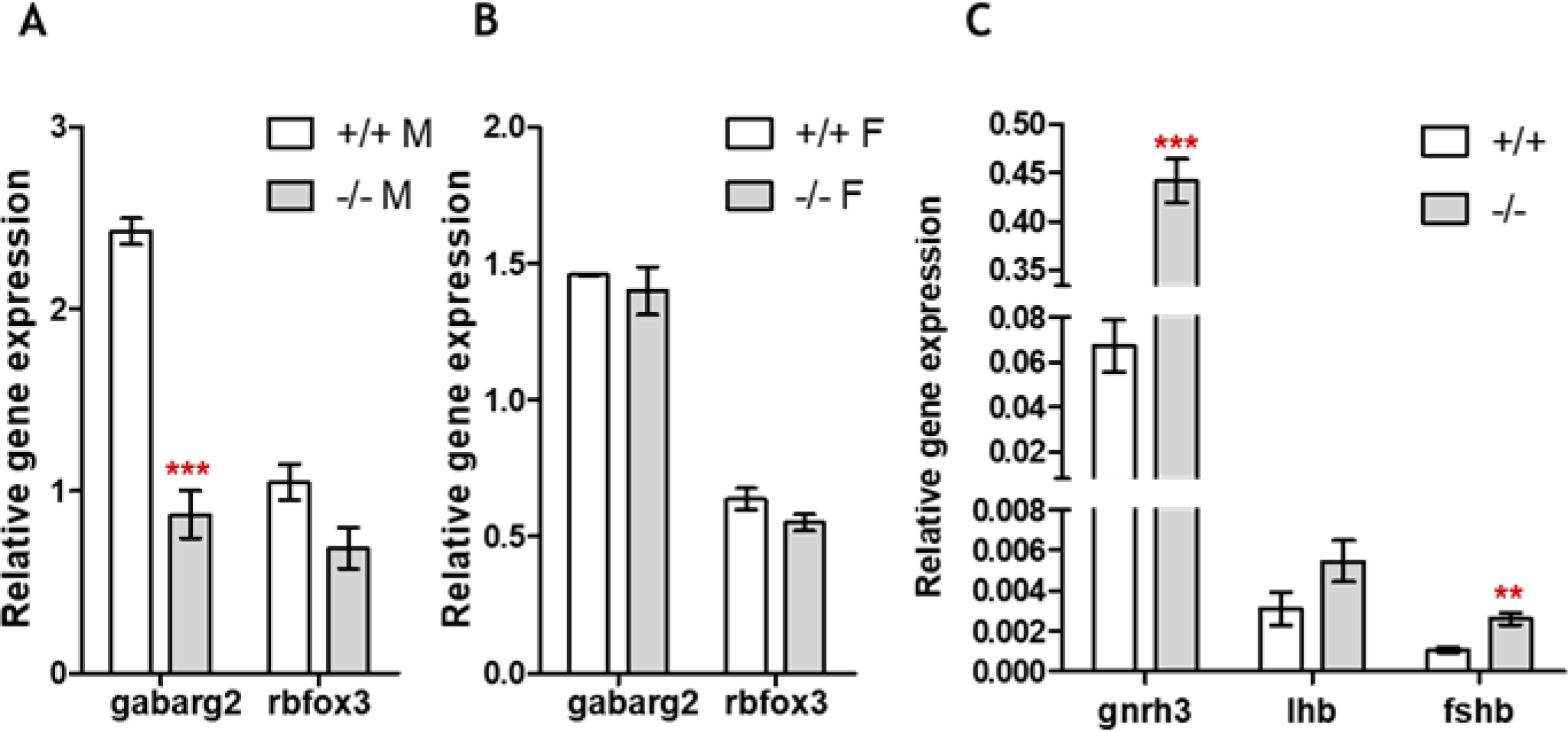
Altered neuroendocrine signalling in *fmr1* KO. A Relative gene expression of *gabarg2* and *rbfox3* in brain tissue from male (A) and female (B) zebrafish. C. Relative expression of *gnrh3*, *fshb* and *lh* mRNA in 5-6mpf KO female brain tissue relative to wildtype (n=3/group).

### Loss of FMRP appears to promote increased oogenesis in juvenile zebrafish

The cytoplasmic polyA element binding proteins (CPEBs), regulate cytoplasmic polyadenylation (CPA) and exert tight spatial and temporal control of translation of maternal mRNAs in the oocyte during oogenesis, oocyte maturation and in the early embryo until zygotic transcription is initiated (Mendez and Richter, 2001). We investigated the impact of FMRP loss on the zebrafish oocyte CPEB *zorba*, the embryonic CPEB and Zorba target, *elrA,* and their targets, using our KO lines. We carried out this analysis in fish of three different ages: juvenile – 1mpf; young adult – 5-6mpf and old adult – 9-12mpf, to cover various stages of ovarian development. We measured the polyadenylation status (considered a direct read-out of translation) using a PAT-PCR assay (polyadenylation tail length PCR assay, Fig S7A) as well as the gene expression profile of the appropriate targets at these time-points.

In the ovaries of 1-month old juvenile fish, where ovary differentiation has just begun and oocytes are in Stage IB, a significant increase in the mRNA levels *zorba* and of several key genes was observed. These included genes involved in oocyte maturation, meiotic entry and progression such as *dazl* (an RNA binding protein that regulates translation, required for germ cell proliferation and germline stem cell (GSC) establishment); *cmos* (a Serine/threonine protein kinase required for oocyte meiosis and microtubule organization); *buc* (the germplasm organiser and homolog of Oskar from Drosophila, regulates the number of germ cells and proper localization of GC genes *nanos, dazl* and *vasa)*; *ccnb1* (the cyclin B1 gene, a component of maturation promotion factor (MPF), required for proper meiotic progression); and RINGO/*spdy,* an atypical activator of cyclin-dependent kinases and necessary for CPEB-directed polyadenylation (Fig. 4A). Overexpression of *dazl* and *buc* are likely to result in excess GC formation, in unexpected locations, mimicking the observations in the *dfmr* KO flies (Bontems et al., 2009; Fukuda et al., 2018). We did not observe a significant difference in the polyA profile of the *zorba* targets at this age, indicating no major changes in translation (Fig 4B).

**Figure 4:**
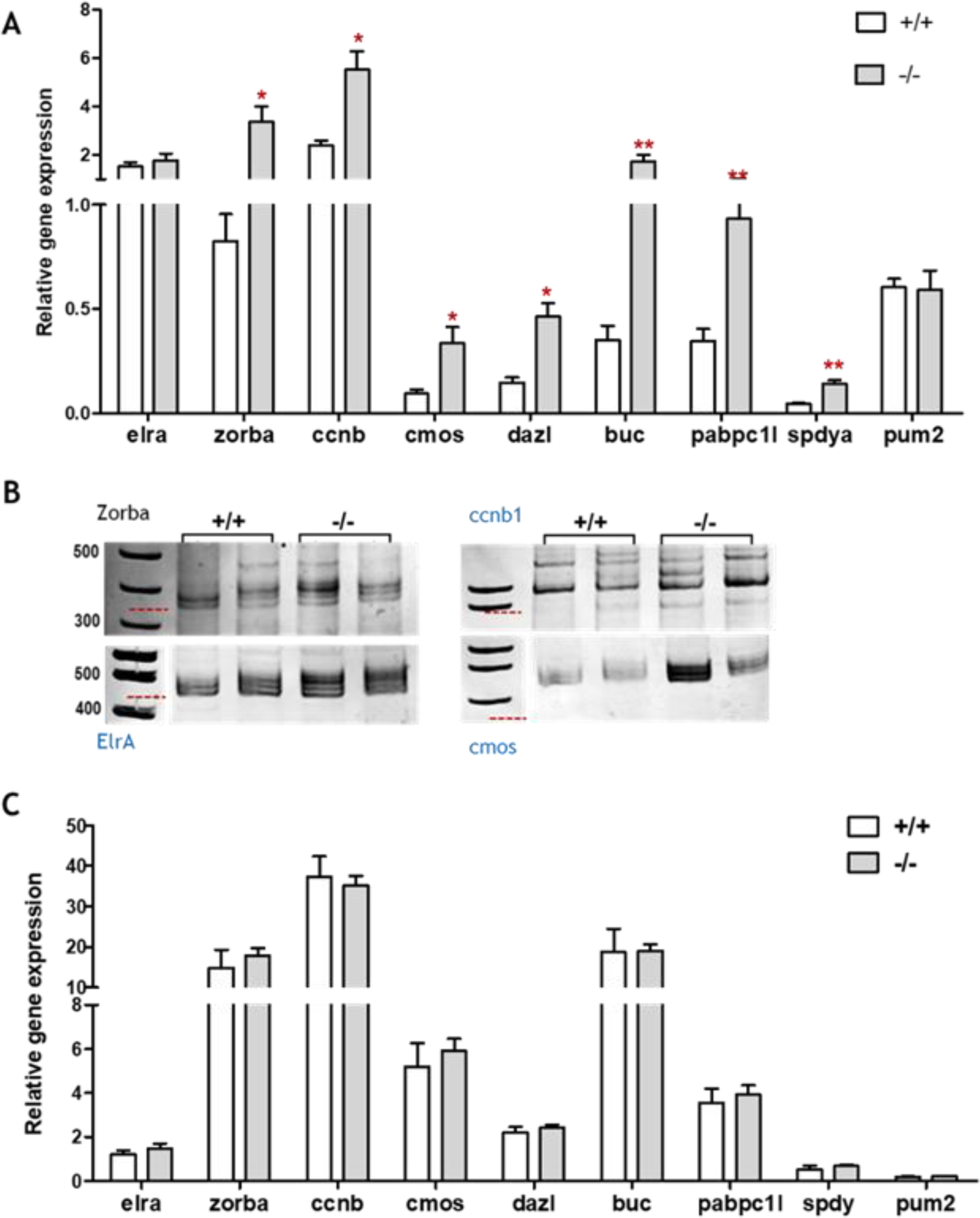
Gene expression profile in juvenile and young adult zebrafish (1mpf and 6mpf): A, C. Relative expression levels of genes involved in CPA, meiotic entry and progression in ovaries (1mpf, A) and oocytes (6mpf, C) n=3 fish/group. B. PAT assay for *zorba, elrA, cmos* and *ccnb1* in the same sample as in A.

Adult zebrafish have asynchronous ovaries, containing follicles (oocytes) of all stages of development, each stage defined according to size and morphology (Clelland and Peng, 2009; Selman et al., 1993). In stage III/IV oocytes from young female fish, the expression of the same markers was neither up nor down regulated in the KOs relative to wildtype, indicating that those oocytes that progressed through maturation are similar to those in wildtype, thus explaining the ability of these fish to produce viable embryos (Fig. 4C).

### Altered oocyte morphology and distribution in apparently normal young knockout females

In order to assess the distribution and morphology of follicles in a young adult female, we undertook histological analysis of ovarian sections in young adult (5-6mpf) females at the same stage in the breeding cycle. H&E stained ovary sections revealed a very different distribution of the different stages of oocytes in the KO, with fewer stage I and more stage III/IV oocytes (representative images of the sections are shown in Fig 5A, B and an assembled image of the ovary cross-section is shown in Fig S11). Additionally, in the KO, a number of large follicles containing fused structures (instead of yolk granules) characteristic of atretic follicles were present, and they lack a well-define zona radiata (Fig 5B and S8B). The mitochondria in these KO oocytes, appeared to be less in number (lower intensity) with a more diffuse architecture as compared to the distinct Balbiani body-like structure observed in wildtype (Fig S8A). While these fish ovulate normally at this stage, their ovaries appear to contain a reduced reserve of good quality oocytes, and resemble the ovary of a much more aged fish.

**Figure 5:**
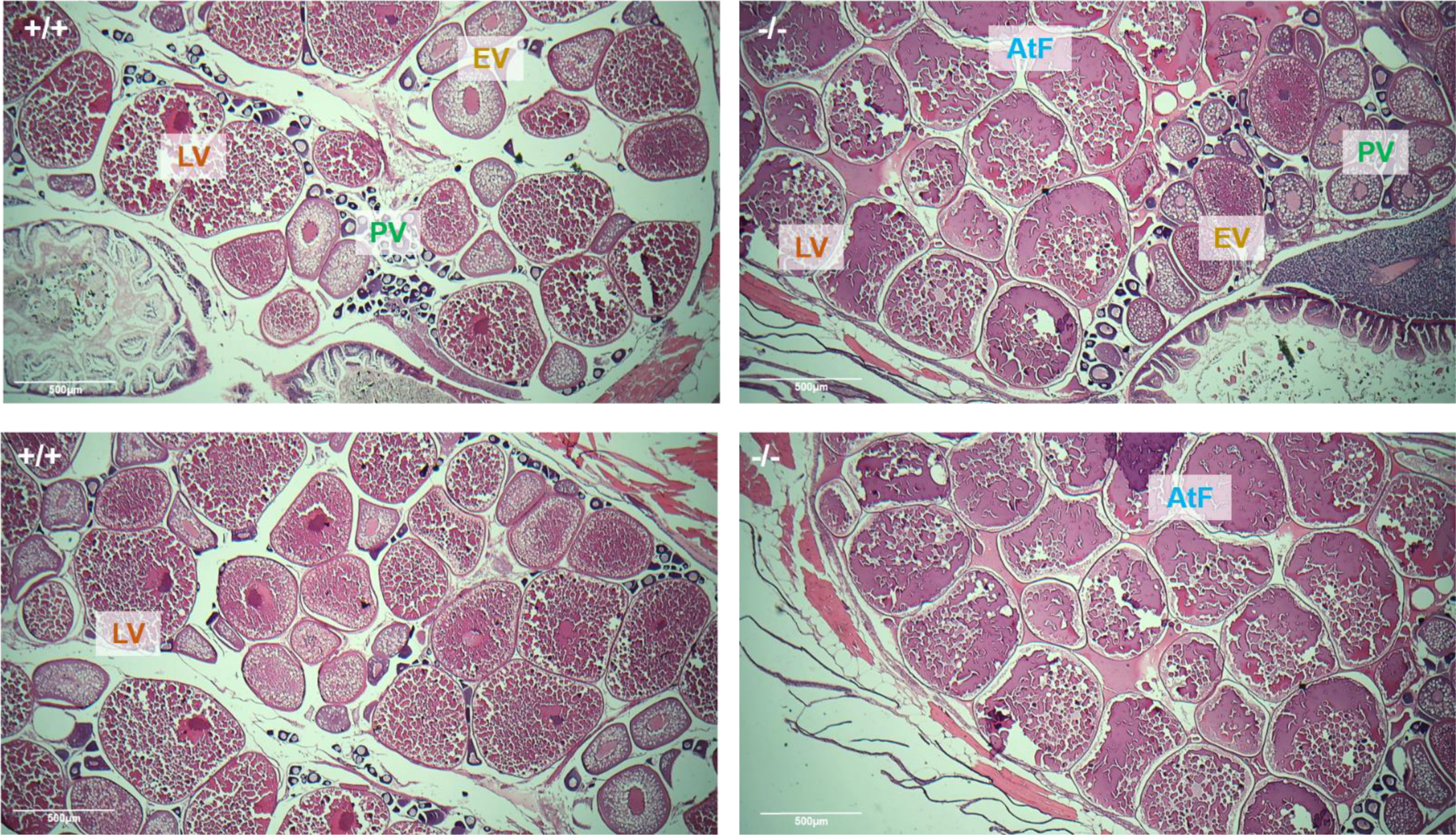
Histological analysis of H&E stained ovarian sections from wildtype (A) and KO (B) zebrafish (5mpf). PV: PreVittelogenic; EV: Early Vittelogenic; LV: Late Vittelogenic; AtF: Atretic follicles.

### Loss of FMRP drastically alters cytoplasmic polyadenylation in late stage oocytes from old females

In oocytes from old FMRP KO females, a drastic reduction in the level of *zorba* mRNA was detected (Fig. 6A, 7A). And a consequent and untimely increase in both the level and polyA tail length of *elrA* mRNA was observed in the KO oocytes (Fig 6A). However, the level and polyA tail length of *elrA* post fertilisation, appear to be similar to that of wildtype (Fig 6A and Fig.S9B). These results are contradictory to the findings from flies, and from juvenile zebrafish ovaries, where FMRP loss results in Zorba upregulation. To validate our findings, we assessed the polyA profile of the mRNA targets of Zorba and ElrA, to see if their polyadenylation was also changed consistently in the oocytes from old KO females. As expected, several targets of Zorba show an altered polyA profile in the KO oocytes, including *buc, ccnb1* and *chtf8* (the orthologue of Drosophila fs(1)K10, a protein required for dorsal-ventral specification and required for egg as well as embryo polarity). Similarly, an ElrA target, *hnrnp1ab*, a protein that in turn controls the translation of several embryonic mRNAs, was also found with increase polyA tail length (Fig. S9C). Unexpectedly, we found a significant decrease in the polyA tail length of an oocyte tubulin gene *tubb4b,* which is not a known target of Zorba or ElrA. Further, we confirmed the drastic decrease in protein levels of Zorba as well as Tubulin, in these oocytes from old females, by IHC as shown in Fig 6C.

**Figure 6:**
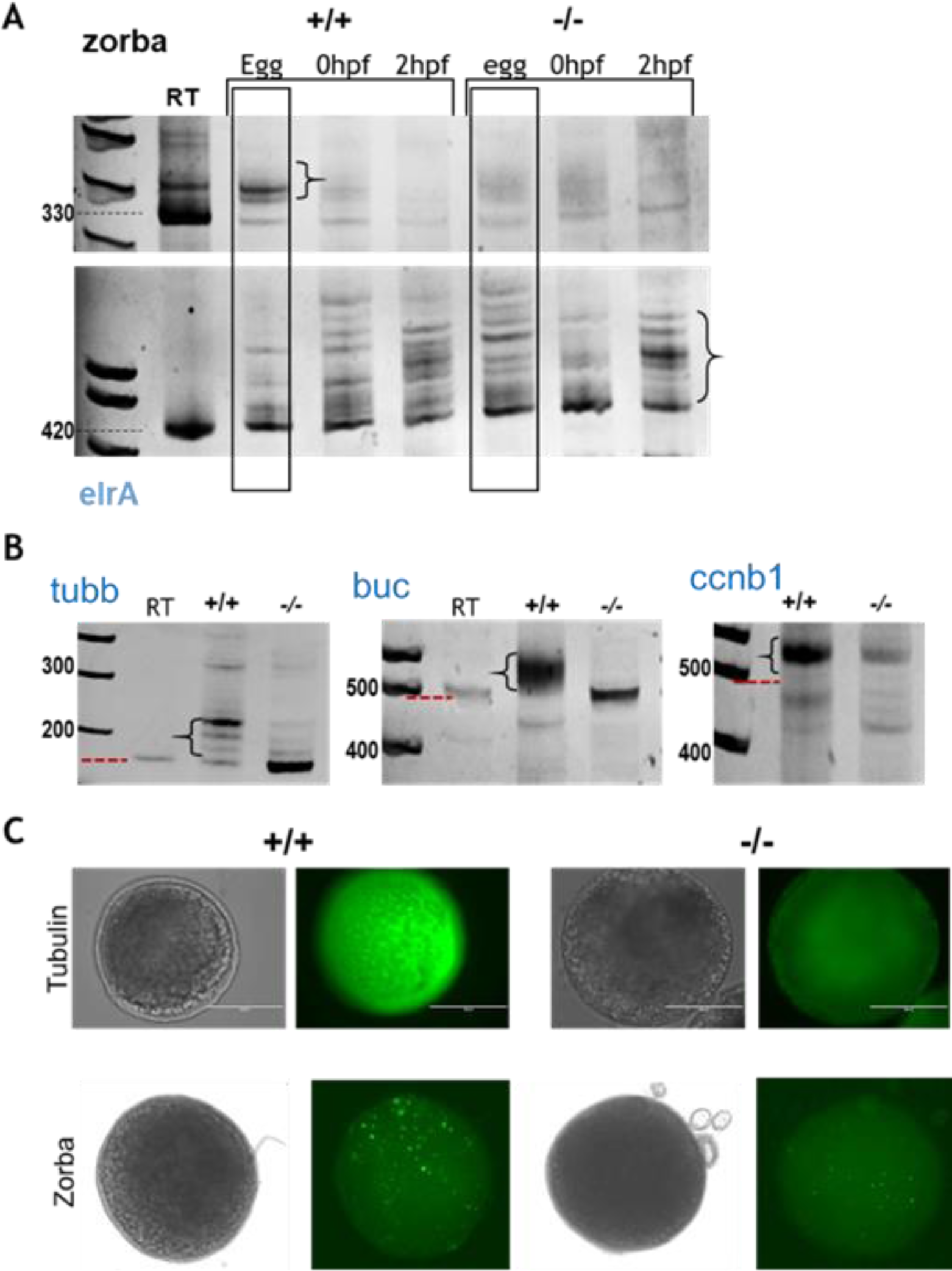
Altered polyadenylation profile of transcripts in oocytes from old knockout fish: **A.** PAT assay for *zorba* and *elrA* from stage III/IV eggs, 0hpf and 2hpf embryos. B. PAT assay in zorba target genes (*buc, ccnb1*) and other transcripts controlled by CPA (*tubb4b* (tubulin)) in the same samples as in A. C. IHC with the α-tubulin and orb (Zorba) antibodies in stage III/IV oocytes from older (9-12mpf) female KO and wildtype fish.

### FMRP loss affects the expression of key genes involved in oocyte maturation

In addition to the polyA changes, a significant decrease in the mRNA levels of several key genes involved in oocyte maturation, meiotic entry and progression such as *dazl*, *cmos*, *buc* and *ccnb1* was also observed (Fig 7A). The magnitude of reduction seen in the FMRP KO oocytes from the 9-12mpf fish is even more than that observed in oocytes from 18mpf wildtype fish (Fig S10A) and hence not just a consequence of aging. Additionally, the FMRP target *pum2* mRNA, an RNA binding translation regulator, was found to be increased (Fig 7B). And the Pum2 mRNA target RINGO/*spdy* which is not regulated by CPA, and is important for meiosis resumption, was also found to be decreased. We measured a drastic decrease in the levels of the embryonic polyA binding protein (ePAB, *pabpc1l)*, the predominant cytoplasmic PABP expressed in oocytes and early embryos which prevents deadenylation of mRNAs (Fig 7B). All of these genes are required for the release from MII and final maturation (Esencan et al., 2019; Ozturk, 2019; Padmanabhan and Richter, 2006). The levels of several tubulin genes expressed in oocytes were also reduced suggesting an impact on the cytoskeleton (Fig 7C). Such drastic changes are likely to severely impact translation and proper localization of mRNAs in the oocyte, thus impacting fertilization and subsequent development of the embryo.

**Figure 7:**
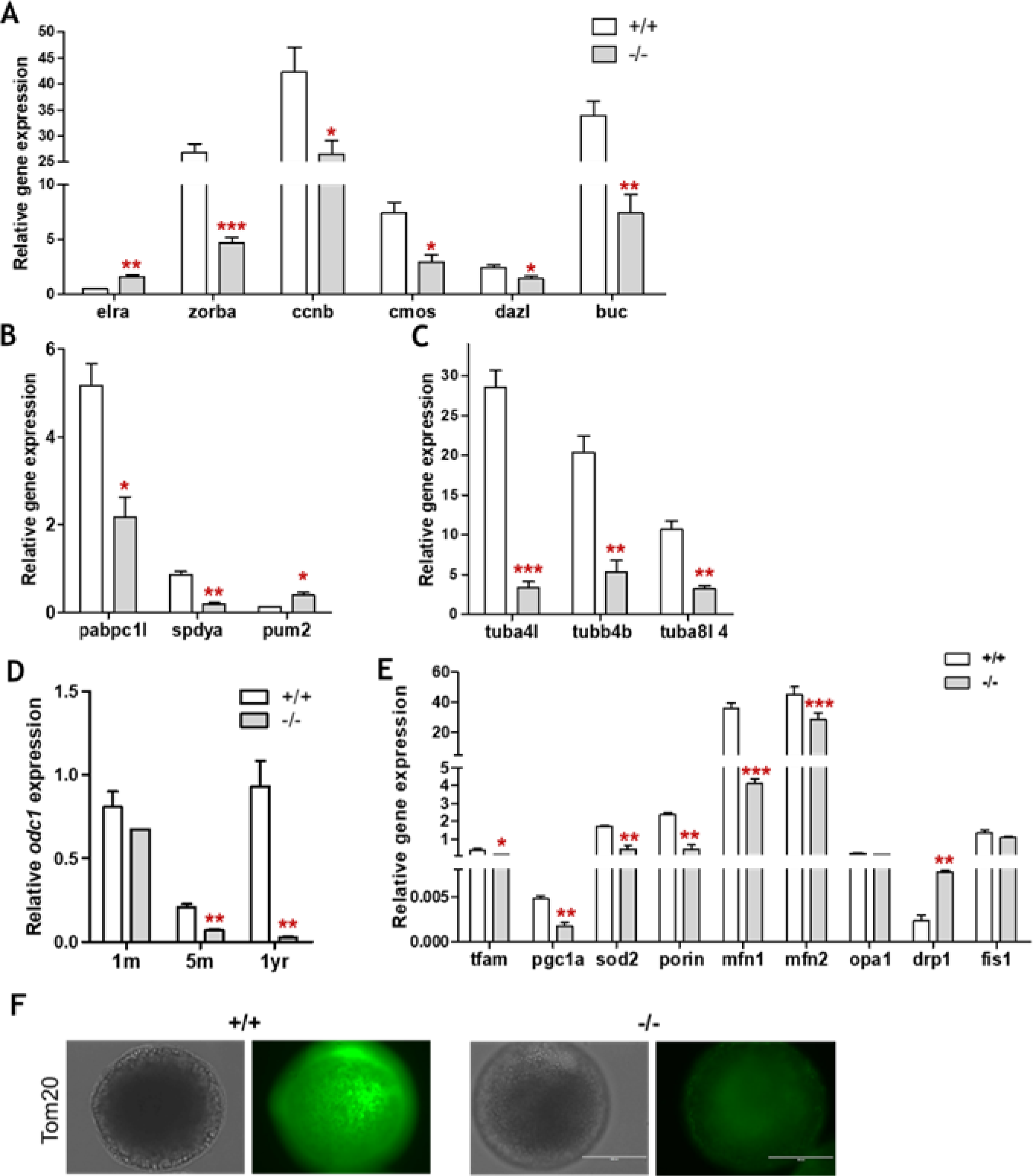
Effect on FMRP loss on gene expression in oocytes from old female zebrafish (9-12mpf): A. Genes involved in PGC specification and differentiation, GSC proliferation, and meiotic entry and progression B. FMRP target *pum2*, its target RINGO/*spdy* and the embryonic PABP. C. three different tubulin genes expressed in oocytes. D. Expression levels of *odc1,* a rate-limiting enzyme in the spermidine synthesis pathway in oocytes from zebrafish of different ages, E. Relative expression of mitochondrial markers. F. IHC with the Tom20 antibody to visualize mitochondrial morphology and intensity in the same oocytes.

To test whether the effect of FMRP loss was the reason for these major changes, we injected an FMRP antibody into stage III/IV oocytes collected from a wildtype female fish. The antibody is expected to sequester FMRP away and mimic the KO. As seen with the KO, the level of *zorba* mRNA was reduced in these oocytes, but not in those injected with a mock antibody (Fig. S10B). This suggests that FMRP is likely to stabilize the *zorba* mRNA and prevent its degradation in Stage III/IV oocytes.

Taken together, these results suggest that FMRP loss affects the transcriptome of oocytes in very different ways in fish of various ages (with oocytes at various stages), with a marked increase in meiotic genes in juveniles, to an unaltered profile in a young adult, all the way to a drastic downregulation of the same markers in an older adult female.

### FMRP loss has a severe impact on mitochondrial health in oocytes

Polyamines have been shown to play an important role in reproductive function, among many other processes. Spermidine levels decrease with age, and directly result in poor fertility in aged female mice (Zhang et al., 2023). We measured the level of the key enzyme involved in spermidine synthesis (*odc1*) and found it to be reduced in the knockouts at all ages (Fig. 6D), and most drastically in the oocytes from older fish. Reduced spermidine results in impaired mitophagy, and increased apoptosis, and may account for the increase in atretic follicles in the KO fish right from 5-6mpf (Fig 5B). In the oocytes from old KO females, we found a decrease in several genes important for mitochondrial biogenesis and function such as *pgc1α, tfam, sod2, porin*; and a significant decrease in the fusion genes *mfn1* and *mfn2* as well as an increase in the fission marker *drp1* (Fig 7E). Further, we observed much weaker and more diffuse signal for mitochondrial staining (IHC with a Tom20 antibody) in oocytes from the KO fish (Fig 7F) suggesting a severe impact of FMRP loss on oocyte mitochondria, which is likely to impact oocyte and subsequently embryo quality.

## Discussion

In this study, we created and characterized *fmr1* KO zebrafish lines, which model certain aspects of human FXS behavior such as hyperactivity and anxiety, in the juvenile and adult stages. These behavior phenotypes are accompanied by and rooted in the altered expression of genes that regulate neuronal function. In addition, we have also catalogued differences in ovarian development and oocyte maturation in the *fmr1* KO females, and report on at least three molecular mechanisms that could contribute to these defects. The first is evidence of increased levels of genes that encode the neuroendocrine hormones that promote oocyte maturation, in the brain of female KO zebrafish. The second is drastic differences in the expression of key genes responsible for PGC specification, GSC proliferation and meiotic progression, and an altered distribution of oocytes at various stages in the KO zebrafish. And finally, we show proof of mitochondrial dysfunction in both the brain and the oocytes of KO zebrafish, which possibly contributes to neuronal cell death, and increased apoptosis and poor quality of oocytes in the older KO females.

FXS has been modeled in zebrafish previously using Morpholino knockdown (Tucker et al., 2006) or a DNAzyme knockdown (Medishetti et al., 2019), a mutant line generated by ENU mutagenesis (den Broeder et al., 2009; Ng et al., 2013) and by CRISPR mediated gene editing (Hu et al., 2020), with each study reporting specific phenotypes. In general, stronger phenotypes were observed with the transient knockdown methods including hyperactivity, altered anxiety, and craniofacial defects in the larvae, with a measurable impact on the mGluR5 signaling cascade. Subtle neuronal phenotypes such as altered brain-wide auditory networks (Constantin et al., 2020), changes in synaptic density and locomotor activity (Shamay-Ramot et al., 2015) and learning and memory deficits (Hu et al., 2020) have been reported in the stable KO lines (hu2787, and the CRISPR mutant) but none report evidence of mGluR5 signaling perturbations. Only a few studies report phenotypes in older fish which include increased anxiety and precocious shoaling at 28dpf (Wu et al., 2017), hyperactivity, anxiolytic-like behavior and impaired learning accompanied by enhanced LTD in adults (Ng et al., 2013). Consistent with the literature, we observe no robust phenotypes in 7dpf larvae, and only minor gene expression changes in olig2, a marker of oligodendrocytes and GFAP, a marker of astrocytes. This is consistent with the hypomyelination observed in infants with FXS (Doll et al., 2021).

However, in older larvae (14dpf) and in adults, we detected robust behavioral changes consistent with increased anxiety, reduced exploration and impaired cognition. FMRP deficient immature neurons in KO mice, have previously been shown to have reduced Huntingtin (*htt*) mRNA and protein levels, which results in impaired mitochondrial function, fragmented mitochondria and defective dendritic maturation (Shen et al., 2019). We found a similar decrease in *htt* mRNA and mitochondrial fusion genes which could be the basis of improper neuronal maturation. FMRP is known to play an important role in astrocytes as well, and neurons cultured on FMRP deficient astrocytes show abnormal dendritic morphology (Jacobs and Doering, 2010). Mitochondrial dysfunction in astrocytes is known to result in reactive gliosis (increased GFAP) and increased neuronal cell death (reduced *rbfox3*) (Fiebig et al., 2019), both of which we observe in the brain from the adult KO zebrafish. These molecular and behavior changes in the adult correlate well with each other, and with changes observed in the mouse model as well as humans with FXS, and have been demonstrated for the first time in the adult FMRP KO zebrafish.

We did not find a convincing rationale for the differences in observed phenotypes in larvae, between the transient knockdown and the stable knockout of FMRP. We or others (Barthelson et al., 2021) found no obvious compensatory gene expression changes. But it is well known that FMRP plays a critical role during early development, where temporal and spatial control of maternal mRNA translation is critical. Therefore, we hypothesized that in the knockout line, compensatory mechanisms at the level of translation could be activated early on to ensure survival through the MZT in the absence of FMRP, which may not continue to be at play in the adult brain or gonads. For example, even though we observed drastic changes in the expression of *zorba* mRNA, and polyadenylation of its targets such as *elrA* and *hnrnp1ab* in the oocytes, we did not observe this to be carried over post fertilization, with both *elrA* and *hnrnp1ab* levels and polyA profile remaining similar to wildtype (Figure 6A and S9C).

FMRP is known to be expressed in the ovaries of flies, mice, rats and humans (Ascano et al., 2012; Takahashi et al., 2015), but there are not many studies delving into the exact function of FMRP during oogenesis. In humans, both males and females with FXS have been reported to be fertile and capable of reproduction (Allingham-Hawkins et al., 1999; Tolmie, 2002). But other than reports suggesting it is possible, there have been no detailed studies on ovarian development, ovarian reserve and the actual window of fertility. Premature ovarian insufficiency (POI) and diminished ovarian reserve (DOR) have been studied extensively in premutation carrier females (PM), of whom 25% are reported to be infertile (Man et al., 2017). It is believed that the increased levels of the toxic *fmr1* RNA in PM carriers is more detrimental to ovarian dysfunction.

The most amount of work has been carried out in Drosophila, where FMRP is implicated in PGC proliferation, GSC specification and in the regulation of translation, especially via the microRNA pathways (Costa et al., 2005; Yang et al., 2007). One report in the mouse Fmr1 KO mouse showed that these mice exhibited an earlier decline in fertility, premature recruitment of follicles due to elevated mTOR activity (Mok-Lin et al., 2018). Our study is the first to investigate reproductive and ovarian phenotypes in FMRP KO zebrafish, and our findings are very similar to that seen in the mouse model, in terms of increased oogenesis in the juvenile stage, premature maturation of follicles in the young adult resulting in the depletion of ovarian reserve, followed by a rapid and early deterioration of oocyte quality in the older females. At a molecular level, we sought to connect the observed phenotypes to the cytoplasmic polyadenylation machinery and its regulation via FMRP. The CPEB1 protein is known to be critical for oogenesis and oocyte maturation in flies, zebrafish and mice (Bally-Cuif et al., 1998; Christerson and Mckearin, 1994; Hake and Richter, 1994; O’Connell et al., 2014; Takahashi et al., 2023). A functional interaction between FMRP and CPEB1 is also well reported in multiple contexts, where both proteins regulate each other’s activities, predominantly in an antagonistic fashion (Oe et al., 2022; Shin et al., 2022). The strongest evidence we see of the FMRP-Zorba functional interaction in our model is in late stage oocytes from older females, where both in the KO and in wildtype oocytes injected with an FMRP antibody, there is a reduction of *zorba* mRNA and protein levels, as well as altered polyA profiles of known Zorba targets (in the KO). Even though this is the opposite of what is seen in the Fmr1 KO mouse brain cortex, and in FMRP KO cell lines, it is possible that this is true in this specific context. These oocytes are of poor quality and resemble those from a much older wildtype female, and studies have demonstrated a decline in both CPEB1 and FMRP levels in aged females, although a causal link has not been established (Takahashi et al., 2015; Takahashi et al., 2023).

The KO fish also show altered expression of genes involved in germline specification, meiotic entry and maturation, explaining the defects seen the distribution of oocytes in various stages as well as the quality of oocytes from older females. The novel finding is the evidence of increased neuroendocrine stimulation in sexually mature KO fish, much like in the mice, which appears to have an additive effect in promoting the precocious maturation of oocytes and the subsequent depletion of the ovarian reserve. However, it remains to be demonstrated in this specific KO, exactly what the downstream implications of this excessive hormonal stimulation are, and if CPEB1 or other RBPs are involved.

Finally, we also see a strong detrimental impact of FMRP loss on mitochondrial function in both oocytes and in the brain, as has been reported in several model systems and contexts. While in the brain, the mitochondrial impairment is due to a reduction in Huntingtin, in oocytes, we connect these findings to a downregulation of the spermidine synthesis pathway. The mechanism by which FMRP regulates this pathway remains to be uncovered.

Our study in the novel *fmr1* KO zebrafish lines expands on the array of neuronal as well as reproductive FXS phenotypes studied in zebrafish, and strongly supports the idea of using this model for identification of new pathways for therapeutic interventions, especially based on the leads identified here. In future studies, it may be interesting and important to determine whether mGluR5 mediated upregulation is present using detailed neurophysiological studies, and thereby reveal alternate pathways if they exist. We also catalogued the reproductive phenotypes in the knockout females, and identified certain changes in gene expression which may be causal. However, detailed studies will be required to identify the all targets of FMRP in oocytes at various stages, and then to map their functional contributions to a successful oocyte maturation program.

## Acknowledgements

We are grateful to Dr.Anil Challa for help with guide design and optimisation of CRISPR protocols in zebrafish, to Kapil K for experimental assistance and to Dr.Pushkar Kulkarni for advice on zebrafish behaviour experiments. We would like to acknowledge funding support from the Department of Biotechnology, Govt. of India [BT/PR28305/GET/119/272/2018]; DBT/Wellcome Trust India Alliance [Clinical Research Center Team Grant (IA/CRC/20/1/600002)] and CSR funding from Dr. Reddy’s Foundation (Center for Rare Disease Models, Dr. Reddy’s Institute of Life Sciences).

## References

Allingham-Hawkins, D. J., Babul-Hirji, R., Chitayat, D., Holden, J. J. A., Yang, K. T., Lee, C., Hudson, R., Gorwill, H., Nolin, S. L., Glicksman, A., et al. (1999). Fragile X premutation is a significant risk factor for premature ovarian failure: The international collaborative POF in fragile X study - Preliminary data. Am. J. Med. Genet. 83,.

Ascano, M., Mukherjee, N., Bandaru, P., Miller, J. B., Nusbaum, J. D., Corcoran, D. L., Langlois, C., Munschauer, M., Dewell, S., Hafner, M., et al. (2012). FMRP targets distinct mRNA sequence elements to regulate protein expression. Nature 492,.

Asiminas, A., Jackson, A. D., Louros, S. R., Till, S. M., Spano, T., Dando, O., Bear, M. F., Chattarji, S., Hardingham, G. E., Osterweil, E. K., et al. (2019). Sustained correction of associative learning deficits after brief, early treatment in a rat model of Fragile X Syndrome. Sci. Transl. Med. 11, 1–11.

Bally-Cuif, L., Schatz, W. J. and Ho, R. K. (1998). Characterization of the zebrafish Orb/CPEB-related RNA-binding protein and localization of maternal components in the zebrafish oocyte. Mech. Dev. 77, 31–47.

Barthelson, K., Baer, L., Dong, Y., Hand, M., Pujic, Z., Newman, M., Goodhill, G. J., Richards, R. I., Pederson, S. M. and Lardelli, M. (2021). Zebrafish Chromosome 14 Gene Differential Expression in the fmr1hu2787 Model of Fragile X Syndrome. Front. Genet. 12, 1–11.

Bear, M. F., Huber, K. M. and Warren, S. T. (2004). The mGluR theory of fragile X mental retardation. Trends Neurosci. 27, 370–377.

Bontems, F., Stein, A., Marlow, F., Lyautey, J., Gupta, T., Mullins, M. C. and Dosch, R. (2009). Bucky Ball Organizes Germ Plasm Assembly in Zebrafish. Curr. Biol. 19, 414–422.

Christerson, L. and Mckearin, D. (1994). orb is required for anteroposterior and dorsoventral patterning during Drosophila oogenesis. Genes Dev. 8 5, 614–28.

Clelland, E. and Peng, C. (2009). Endocrine/paracrine control of zebrafish ovarian development. Mol. Cell. Endocrinol. 312,.

Clinical Synopsis - #300624 - FRAGILE X SYNDROME; FXS - OMIM.

Constantin, L., Poulsen, R. E., Scholz, L. A., Favre-Bulle, I. A., Taylor, M. A., Sun, B., Goodhill, G. J., Vanwalleghem, G. C. and Scott, E. K. (2020). Altered brain-wide auditory networks in a zebrafish model of fragile X syndrome. BMC Biol. 18, 125.

Costa, A., Wang, Y., Dockendorff, T. C., Erdjument-Bromage, H., Tempst, P., Schedl, P. and Jongens, T. (2005). The Drosophila fragile X protein functions as a negative regulator in the orb autoregulatory pathway. Dev. Cell 8 3, 331–42.

den Broeder, M. J., van der Linde, H., Brouwer, J. R., Oostra, B. a, Willemsen, R. and Ketting, R. F. (2009). Generation and characterization of FMR1 knockout zebrafish. PLoS One 4, e7910.

Deshpande, G., Calhoun, G. and Schedl, P. (2006). The drosophila fragile X protein dFMR1 is required during early embryogenesis for pole cell formation and rapid nuclear division cycles. Genetics 174, 1287–1298.

Doll, C. A., Scott, K. and Appel, B. (2021). Fmrp regulates oligodendrocyte lineage cell specification and differentiation. Glia 69, 2349–2361.

Epstein, A. M., Bauer, C. R., Ho, A., Bosco, G. and Zarnescu, D. (2009). Drosophila Fragile X protein controls cellular proliferation by regulating cbl levels in the ovary. Dev. Biol. 330 1, 83– 92.

Esencan, E., Kallen, A., Zhang, M. and Seli, E. (2019). Translational activation of maternally derived mRNAs in oocytes and early embryos and the role of embryonic poly(A) binding protein (EPAB). Biol. Reprod. 100, 1147–1157.

Fiebig, C., Keiner, S., Ebert, B., Schäffner, I., Jagasia, R., Lie, D. C. and Beckervordersandforth, R. (2019). Mitochondrial dysfunction in astrocytes impairs the generation of reactive astrocytes and enhances neuronal cell death in the cortex upon photothrombotic lesion. Front. Mol. Neurosci. 12,.

Fukuda, K., Masuda, A., Naka, T., Suzuki, A., Kato, Y. and Saga, Y. (2018). Requirement of the 3′-UTR-dependent suppression of DAZL in oocytes for pre-implantation mouse development. PLoS Genet. 14,.

Hake, L. and Richter, J. (1994). CPEB is a specificity factor that mediates cytoplasmic polyadenylation during Xenopus oocyte maturation. Cell 79, 617–627.

Hammond-Weinberger, D. R. and Zeruth, G. T. (2019). Whole mount immunohistochemistry in zebrafish embryos and larvae. J. Vis. Exp. 2020,.

Hu, J., Chen, L., Yin, J., Yin, H., Huang, Y. and Tian, J. (2020). Hyperactivity, Memory Defects, and Craniofacial Abnormalities in Zebrafish fmr1 Mutant Larvae. Behav. Genet.

Jacobs, S. and Doering, L. C. (2010). Astrocytes prevent abnormal neuronal development in the fragile X mouse. J. Neurosci. 30,.

Kalueff, A. V. (2017). The rights and wrongs of Zebrafish: Behavioral phenotyping of zebrafish. (ed. Kalueff, A. V.) Springer.

Kalueff, A. V, Gebhardt, M., Stewart, A. M., Cachat, J. M., Brimmer, M., Chawla, J. S., Craddock, C., Kyzar, E. J., Roth, A., Landsman, S., et al. (2013). Towards a comprehensive catalog of zebrafish behavior 1.0 and beyond. Zebrafish 10, 70–86.

Krisher, R. L. (2004). The effect of oocyte quality on development. J. Anim. Sci. 82 E-Suppl,.

Malecki, C., Hambly, B. D., Jeremy, R. W. and Robertson, E. N. (2020). The RNA-binding fragile-X mental retardation protein and its role beyond the brain. Biophys. Rev. 12,.

Man, L., Lekovich, J., Rosenwaks, Z. and Gerhardt, J. (2017). Fragile X-associated diminished ovarian reserve and primary ovarian insufficiency from molecular mechanisms to clinical manifestations. Front. Mol. Neurosci. 10,.

Medishetti, R., Rani, R., Kavati, S., Mahilkar, A., Akella, V., Saxena, U., Kulkarni, P. and Sevilimedu, A. (2019). A DNAzyme based knockdown model for Fragile-X syndrome in zebrafish reveals a critical window for therapeutic intervention. J. Pharmacol. Toxicol. Methods 101, 106656.

Medishetti, R., Balamurugan, K., Yadavalli, K., Rani, R., Sevilimedu, A., Challa, A. K., Parsa, K. and Chatti, K. (2022). CRISPR-Cas9-induced gene knockout in zebrafish. STAR Protoc. 3, 101779.

Mendez, R. and Richter, J. D. (2001). Translational control by CPEB: A means to the end. Nat. Rev. Mol. Cell Biol. 2, 521–529.

Mok-Lin, E., Ascano, M., Serganov, A., Rosenwaks, Z., Tuschl, T. and Williams, Z. (2018). Premature recruitment of oocyte pool and increased mTOR activity in Fmr1 knockout mice and reversal of phenotype with rapamycin. Sci. Rep. 8,.

Monzo, K., Dowd, S. R., Minden, J. S. and Sisson, J. C. (2010). Proteomic analysis reveals CCT is a target of Fragile X mental retardation protein regulation in Drosophila. Dev. Biol. 340, 408– 418.

Ng, M.-C., Yang, Y.-L. and Lu, K.-T. (2013). Behavioral and synaptic circuit features in a zebrafish model of fragile X syndrome. PLoS One 8, e51456.

O’Connell, M. L., Cavallo, W. C. and Firnberg, M. (2014). The expression of CPEB proteins is sequentially regulated during zebrafish oogenesis and embryogenesis. Mol. Reprod. Dev. 81, 376–387.

Oe, S., Hayashi, S., Tanaka, S., Koike, T., Hirahara, Y., Seki-Omura, R., Kakizaki, R., Sakamoto, S., Nakano, Y., Noda, Y., et al. (2022). Cytoplasmic Polyadenylation Element-Binding Protein 1 Post-transcriptionally Regulates Fragile X Mental Retardation 1 Expression Through 3′ Untranslated Region in Central Nervous System Neurons. Front. Cell. Neurosci. 16, 1–13.

Ozturk, S. (2019). The translational functions of embryonic poly(A)-binding protein during gametogenesis and early embryo development. Mol. Reprod. Dev. 86, 1548–1560.

Padmanabhan, K. and Richter, J. D. (2006). Regulated Pumilio-2 binding controls RINGO/Spy mRNA translation and CPEB activation. Genes Dev. 20,.

Richter, J. D. and Zhao, X. (2021). The molecular biology of FMRP: new insights into fragile X syndrome. Nat. Rev. Neurosci. 22,.

Rosario, R., Childs, A. J. and Anderson, R. A. (2017). RNA-binding proteins in human oogenesis: Balancing differentiation and self-renewal in the female fetal germline. Stem Cell Res. 21,.

Salcedo-Arellano, M. J., Dufour, B., McLennan, Y., Martinez-Cerdeno, V. and Hagerman, R. (2020). Fragile X syndrome and associated disorders: Clinical aspects and pathology. Neurobiol. Dis. 136, 104740.

Sallés, F. J. and Strickland, S. (1995). Rapid and sensitive analysis of mRNA polyadenylation states by PCR. Genome Res. 4, 317–321.

Santoro, M. R., Bray, S. M. and Warren, S. T. (2012). Molecular Mechanisms of Fragile X Syndrome: A Twenty-Year Perspective. Annu. Rev. Pathol. Mech. Dis. 7, 219–245.

Selman, K., Wallace, R. A., Sarka, A. and Qi, X. (1993). Stages of oocyte development in the zebrafish, Brachydanio rerio. J. Morphol. 218,.

Seong-Kyu, H. A. N., Abraham, I. M. and Herbison, A. E. (2002). Effect of GABA on GnRH neurons switches from depolarization to hyperpolarization at puberty in the female mouse. Endocrinology 143,.

Shamay-Ramot, A., Khermesh, K., Porath, H. T., Barak, M., Pinto, Y., Wachtel, C., Zilberberg, A., Lerer-Goldshtein, T., Efroni, S., Levanon, E. Y., et al. (2015). Fmrp Interacts with Adar and Regulates RNA Editing, Synaptic Density and Locomotor Activity in Zebrafish. PLOS Genet. 11, e1005702.

Shen, M., Wang, F., Li, M., Sah, N., Stockton, M. E., Tidei, J. J., Gao, Y., Korabelnikov, T., Kannan, S., Vevea, J. D., et al. (2019). Reduced mitochondrial fusion and Huntingtin levels contribute to impaired dendritic maturation and behavioral deficits in Fmr1-mutant mice. Nat. Neurosci. 22, 386–400.

Shin, J., Paek, K. Y., Chikhaoui, L., Jung, S., Ponny, S., Suzuki, Y., Padmanabhan, K. and Richter, J. D. (2022). Oppositional poly(A) tail length regulation by FMRP and CPEB1. Rna 28, 756–765.

Takahashi, N., Tarumi, W., Itoh, M. T. and Ishizuka, B. (2015). The Stage-And Cell Type-Specific Localization of Fragile X Mental Retardation Protein in Rat Ovaries. Reprod. Sci. 22, 1524–1529.

Takahashi, N., Franciosi, F., Daldello, E. M., Luong, X. G., Althoff, P., Wang, X. and Conti, M. (2023). CPEB1-dependent disruption of the mRNA translation program in oocytes during maternal aging. Nat. Commun. 14,.

Tolmie, J. (2002). Fragile X Syndrome - Diagnosis, Treatment and Research: Third edition. Editors Randi Jenssen Hagerman, Paul J Hagerman. pound65.50 HB, pound31.00 PB. Baltimore: The Johns Hopkins University Press. 2002. ISBN 0-8018-6844-0. J. Med. Genet. 39,.

Tucker, B., Richards, R. and Lardelli, M. (2004). Expression of three zebrafish orthologs of human FMR1-related genes and their phylogenetic relationships. Dev. Genes Evol. 214, 567–74.

Tucker, B., Richards, R. I. and Lardelli, M. (2006). Contribution of mGluR and Fmr1 functional pathways to neurite morphogenesis, craniofacial development and fragile X syndrome. 15, 3446–3458.

Villa, P. A., Lainez, N. M., Jonak, C. R., Berlin, S. C., Ethell, I. M. and Coss, D. (2023). Altered GnRH neuron and ovarian innervation characterize reproductive dysfunction linked to the Fragile X messenger ribonucleoprotein (Fmr1) gene mutation. Front. Endocrinol. (Lausanne). 14, 1–19.

Watanabe, M., Fukuda, A. and Nabekura, J. (2014). The role of GABA in the regulation of GnRH neurons. Front. Neurosci. 8,.

Wierson, W. A., Welker, J. M., Almeida, M. P., Mann, C. M., Webster, D. A., Torrie, M. E., Weiss, T. J., Kambakam, S., Vollbrecht, M. K., Lan, M., et al. (2020). Efficient targeted integration directed by short homology in zebrafish and mammalian cells. Elife 9, 1–25.

Wu, Y.-J., Hsu, M.-T., Ng, M.-C., Amstislavskaya, T. G., Tikhonova, M. A., Yang, Y.-L. and Lu, K.-T. (2017). Fragile X Mental Retardation-1 Knockout Zebrafish Shows Precocious Development in Social Behavior. Zebrafish 14, 438–443.

Yang, L., Duan, R., Chen, D., Wang, J., Chen, D. and Jin, P. (2007). Fragile X mental retardation protein modulates the fate of germline stem cells in Drosophila. Hum. Mol. Genet. 16,.

Zhang, Y., Bai, J., Cui, Z., Li, Y., Gao, Q., Miao, Y. and Xiong, B. (2023). Polyamine metabolite spermidine rejuvenates oocyte quality by enhancing mitophagy during female reproductive aging. Nat. Aging 3, 1372–1386.

