## Supplementary Information for "Impaired ovarian development in a zebrafish *fmr1* knockout model"

[1] Center for Innovation in Molecular and Pharmaceutical Sciences, Dr. Reddy's Institute of Life Sciences, University of Hyderabad Campus, Gachibowli, Hyderabad, Telangana,500046, India. Tel: +91-40-66571500

**Supplementary Figure and Tables:**

**Supplementary Table 1:** List of primers and oligonucleotides used in this study

**Supplementary Table 2:** List of antibodies used in this study

**Supplementary Table 3:** Fold change in gene expression of FMRP target genes in brain tissue from KO fish with respect to wildtype fish (n=3).

**Supplementary Figure 1: A**. In vitro catalytic cleavage assay with the synthesized sgRNA-Cas9 RNPs containing sg1, sg2 or both, incubated with the target amplicon (C) containing the guide cut sites. Lower sized bands are cleavage products. **B.** HMA-PCR of F1 progeny from the outcross showing the pattern for each of the two types of edited alleles. **C**. Predicted protein sequence of the mutant *fmr1* allele surrounding the guide cut site.

**Supplementary Figure 2: A.** Fold change in the expression levels of indicated mRNAs in 7dpf KO larvae as compared to wildtype (n=3) **B.** Measurement of FMRP levels by Western blot in brain tissue from adult KO and wildtype zebrafish.

**Supplementary Figure 3:** A. Representative images of 6dpf larvae stained with Alcian Blue to observe craniofacial structures. B-H Quantification of various parameters from the Alcian blue stained images (n=6/group).

**Supplementary Figure 4: A, B**. Locomotion tracking in individual larvae (7 and 10dpf) in the Zebrabox automated locomotion tracking system. **C**. Average distance moved in the last minute with light (Last L), and the first minute in the dark (First D) during the light-to dark transitions showing increased slope of the KO larvae transition **D**. Average distance moved in the last minute in dark (Last D), and the first minute in with light (First L) during the dark-to-light transitions showing increased deceleration of the KO larvae transition. C and D just a stronger response to either transition in the KO larvae as compared to wildtype.

**Supplementary Figure 5: A.** Fold change the in mRNA expression of indicated genes at 7dpf in KO versus wildtype. **B.** Measurement of the mGluR5 protein and the downstream signalling cascade (p-ERK, p-AKT) in 7dpf KO and wildtype larvae.

**Supplementary Figure 6:** Measurement of the levels of various proteins in the brain tissue of adult zebrafish, by Western blot **A**. mGluR5, **B** p-AKT and AKT, **C** p-ERK and ERK, **D** Mfn2, **E** Opa1 and **F** Zorba

**Supplementary Figure 7:** A schematic illustrating the principle of the PAT-PCR assay. This assay measures the length of the polyA tail on a specific gene transcript. First phospho-oligodT and an anchor reverse primer containing an oligodT stretch, are annealed to the mRNA. The phospho-oligos are ligated using T4 DNA ligase, and RT enzyme is added to generate the cDNA. Subsequently, PCR is performed using a gene specific forward primer in the 3'UTR, and the anchor dT as reverse primer.

**Supplementary Figure 8: A.** Representative images of Tom20 IHC to reveal the structure and number of mitochondria in the KO and wildtype oocytes**. B.** Bright field images of the stage III/IV (0.4-0.7mm) oocytes from KO and wildtype fish (9mpf). The KO oocytes do not appear to have a defined chorion (or ZR)

**Supplementary Figure 9** Relative expression levels of *zorba* **(A)** and *elrA* **(B)** in oocytes and embryos from wildtype and KO fish (9-12mpf, n=3). **C.**PAT PCR with hnrnp1ab, a target of ElrA in oocytes, 0 and 2hpf embryos. RT: Normal RT-PCR is performed using anchordT as the reverse primer to illustrate the size of the amplicon without polyA.

**Supplementary Figure 10: A.** Expression levels of representative genes in wildtype (9-12mpf), aged wildtype (18mpf) and knockout (9-12mpf) fish to illustrate that aging alone does not explain the drastic decrease in these specific genes in the KO oocytes. The reduction is dependent on FMRP loss. **B.** Measurement of *zorba* mRNA levels in control oocytes, oocytes injected with a mock antibody, and those injected with FMRP antibody. Quantification of n=3 experiments is shown on the right.

**Supplementary Figure 11:** Image of the cross section of the entire ovary from wildtype (+/+) and KO (-/-) fish, assembled from multiple tiled images obtained.

**Table S1**

| NAME | OLIGONUCLEOTIDE SEQUENCE |
|---|---|
| DR_FMR1EX3-C1 | TAATACGACTCACTATAG GAGGTGGAAACCGAACATCC GTTTTAGAGCTAGAA |
| DR_FMR1EX4-C1 | TAATACGACTCACTATAG GTACTCTAGGGCTAATGATA GTTTTAGAGCTAGAA |
| TAIL OLIGO | AAAAGCACCGACTCGGTGCCACTTTTTCAAGTTGATAACGGACTAGCCTTATTTTAACTTG CTATTTCTAGCTCTAAAAC |
| T7 PROMOTER FP | GTAATACGACTCACTATA |
| TAIL OILGO RP | AAAAGCACCGACTCGGTG |
| FMR GENOTYPING FP | CTCACTTCAATTTCAGATGCATACAATG |
| FMR GENOTYPING RP | CTTCTCCTTTCACCATGCGAAC |
| FMRQPCR FP | AGCGAGTGGAGCTTTCTA |
| FMRQPCR RP | CCTTTCACCATGCGAACT |
| RNAPD FP | CCAGATTCAGCCGCTTCAAG |
| RNAPD RP | CAAACTGGGAATGAGGGCTT |
| EGR1-FP | TCTACCAGTCTCAGCTCATC |
| EGR1 RP | CTGTGTGTGTGCGAATGT |
| HOMER1A FP | GAGTGTGTGTGTGTGTGT |
| HOMER1A RP | GTGTGCTGTCGTAGAAGTAG |
| GFAP FP | TTGTGCGAACTGTTGAGACC |
| GFAP RP | AGCAGGGAAAGTTGGTGAGA |
| OLIG2 FP | TTGCACCTGCTACCGGGAAT |
| OLIG2 RP | CTTGACGGCGGACAGAAAAAG |
| AQP4 FP | CAGTGTTTCGGGCACATCA |
| AQP4 RP | ATGCACACAGACAGACCAATAG |
| FXR1S FP | CCTCGTTACTGTGGCCGATTA |
| FXR1S RP | CGTTGCTCACAGATTCAGCAG |
| FXR1L FP | TCGATGGAGCTGAAGCCAAA |
| FXR1L RP | AGCTCGCGATATGTAATCGGC |
| FXR2 FP | AAGCGAAAATGGACTGGAAGAG |
| FXR2 RP | AACAGTAACTGGCTGTCGGTCA |
| BDNF FP | TGCAAGTGTTGGTCTTGGAG |
| BDNF RP | CAGCTCTCATGCAACTGAAG |
| FOSL2 FP | GTACCAGGATTACACCGGGA |
| FOSL3 RP | CAGGCATGTCTATTCGGTAC |
| MTOR FP | CCCAGACTTATTCGCCCATAC |
| MTOR RP | CCATTTCCTCATCTCCAGTCC |
| SOX9A FP | CAGCATGGGAGAAGTGCACT |
| SOX9A RP | GACGTCCGGTGTTTTCCTGA |
| ELRA PAT FP | GTTTCTCCCCGTCAGAGGTG |
| TUBB4 PAT FP | CCAGTCTGTTCCATCTTGTG |
| ZORBA PAT FP | TCAGCAGGTGTTGTGGGTGTGA |
| ANCHORT | GCGAGCTCCGCGGCCGCGTTTTTTTTTTTTT |
| P-OILGO DT | TTTTTTTTTTTTTTTTTT |
| ZORBA RTPCR RP | GACAGAGGAGCTTGGGAAATAGC |

| | |
|---|---|
| ELRA RTPCR RP | GCAAACCATAAACAGAAGATGCTGATGC |
| NRXN 1 FP | GAGCAGTAGCGATGAGATTAC |
| NRXN 1 RP | ACTACCGCCGACATAGAA |
| NGLN 3 FP | CTGCTGACTCTTTCCCATTAT |
| NGLN 3 RP | CCTGCTCCACTAGTTCTTTG |
| SHANK 3A FP | GAGGTAGAGGAGGAGGATTT |
| SHANK 3A RP | CATGGGAGGATGTAGTTTAC |
| DLGAP1A PAT FP | AAGCTCACCCTCCTCTGGCC |
| DLGAP1A RP | GATGCTGTCTGCGCTCTCGG |
| DLGAP1B PAT FP | GAGAGGAAGGAGAGGCGCGT |
| DLGAP1B RP | TTCGATGCTGTCGGCGCATT |
| SURF1 PAT FP | TGTGGAGGCAAACAGATGGCA |
| SURF1 RP | TGCCCTCCTATTGGCCCACC |
| SIRT1 PAT FP | AAGTTCCGTGAGTGCCCCTG |
| SIRT1 RP | ACAACTCGTAATATGCATGT |
| CAMTA1A PAT FP | CCCAGAAAATAGCGGCCCTGT |
| CAMTA1A RP | TGCATGGCTTCAGCAGCTGA |
| CAMTA1B PAT FP | TCGGCCGTCTCCCTCTTTCT |
| CAMTA1B RP | TGCATGGCTTCAGCAGCTGA |
| BUC PAT FP | CACCTTGCTGCATTTCCAGTGCT |
| BUC FP | TCCACCAGCAAAGGCCAAGAA |
| BUC RP | TCACCCCTTGAACCAGCTGC |
| CCNB1 PAT FP | TCACTGCCATGGGTTGAGCA |
| CCNB1 FP | GTGAGGGTCAACGAGGGCCT |
| CCNB1 RP | TGCTCAACCCATGGCAGTGA |
| MFN1 FP | CTCGGTCCCGTCAACGCCAA |
| MFN1 RP | ACTGAACCACCGCTGGGGCT |
| MFN2 FP | GCTGGGACGCATCGGCCAAT |
| MFN2 RP | GAGCGATCCACCACCCGCAG |
| OPA1 FP | GCCGGAAGTGTAGTTACCTG |
| OPA1 RP | AGGTGGTCTCTGTGGGTTGT |
| HTT FP | ATGGCCATGTGGAGCCTGTCCT |
| HTT RP | ACGGACTACCAGGGGAAGCCAC |
| DRP1 FP | GAGGAGCAGAGACGTAACCG |
| DRP1 RP | CGGTTGATCAGTGGGTGACA |
| TFAM FP | GGGAACTACTACGCAGGCAA |
| TFAM RP | TGTGCTCCTCCCACGATTTG |
| TUBA41 RP | TCATCCTCCCCGCAGTCATCAG |
| TUBB4B FP | CAGGCAGTTACCATGGCGACAG |
| TUBB4B RP | CCCTTGGCCCAGTTGTTTCCAG |
| TUBA 814 PAT FP | ATGAAGAGGTGGGCACAGAC |
| TUBA 814 QPCR RP | TGCTAGGTTGTGCAAGGGTC |
| MOS FP | TGTACGCTGTTGTGGCGTAT |
| MOS RP | GACGAGCTTATCTGCGGTCG |
| DAZL FP | GCCCCACCATCACCTGTATT |
| DAZL RP | AAGCCTCGTCCTTTGCTCTC |
| PABPC1L FP | GAAACAACTACTGGGCGAGC |

| | |
|---|---|
| PABPC1L RP | GCTACTGCCTCTTCGACCTT |
| SPDYA FP | TCACACAGAGCCAGCAAGAC |
| SPDYA RP | TGATCTGCCTGATGATGGAGC |
| PUM2 FP | CTCTACACCATGATGAAGGACC |
| PUM2 RP | CAATAGGGCCTAGATCTGATCC |
| RBFOX3 FP | TGGTACAGACAGACGGATCT |
| RBFOX3 RP | GGGAATGTTGGAGACGTGTAA |
| FIS1 FP | GAACTGGTTCACACAAGCAA |
| FIS1 RP | GACACGGCCAAACCAATAAG |
| ODC1 FP | TGTCCAATGACCGAACCCTG |
| ODC1 RP | ATACTGCAGGGGTACATGCG |
| PGC1A FP | GGGCAAACAGCCAATTGAGG |
| PGC1A RP | TTTCGTCCATGTTGCGAGGA |
| FSHB FP | GCAGCTGTCGACTCACAAAC |
| FSHB RP | CACGGGGTACACGAAGACTG |
| LHB FP | GCTGTCCAAAATGCCTGGTG |
| LHB RP | CAGTCGGGCAGGTTAATGGT |
| GABARG2 FP | CGGCTATGGACCTCTTCGTG |
| GABARG2 RP | GAGCCGCAGGAGAGGATTTT |
| GNRH3FP | GGCTTCCCGGTGGAAAAAGA |
| GNRH3RP | TCCCCGTCTGTCTGGAAATC |
| MNSOD(SOD2) FP | AAGTCTCCCTTCAGCCTGCATT |
| MNSOD(SOD2) RP | GCTTTATGGCCTCCAACAGCTC |
| PORIN(VDAC1) FP | TGTGGGATACAAGACCGACGAG |
| PORIN(VDAC1) RP | GCGTCGCTGTCGATTTGGTATT |

**Table S2:**

| S.No | Name | Company, Catalog # | Dilution used |
|---|---|---|---|
| 1 | GFAP | DSHB, 8E17 | 1:1000 (WB), |
| 2 | Phospho-p44/42 MAPK (Erk1/2) (Thr202/Tyr204) | CST (9101) | 1:5000 (WB) |
| 3 | Erk | CST (9102) | 1:5000 (WB) |
| 4 | Akt | CST (9272) | 1:5000 (WB) |
| 5 | Phospho-Akt (SER473)(D9E) XP | CST (4060) | 1:5000 (WB) |
| 6 | α-Tubulin | sigma (T6199) | 1:5000 (WB), 1:200 (IHC) |
| 7 | mGluR5 | Merck (ab5675) | 1:5000 (WB) |
| 8 | Fmr1 | Genetex (GTX125996) | 1:5000 (WB) |
| 9 | Orb | DSHB (orb 6H4, AB528419) | 1:1000 (WB), 1:50 (IHC) |
| 10 | Tom20 | Proteintech (11802-1-AP) | 1:2000 (WB), 1:200 (IHC) |
| 11 | Opa1 | CST(80471) | 1:2000 (WB) |
| 12 | Mfn2 | CST (9482) | 1:2000 (WB) |

**Table S3**

| | Description | Category | Average FC | SEM | Signifi-cance |
|---|---|---|---|---|---|
| mtor | Mechanistic target of rapamycin (serine/threonine kinase) | FMRP CLIP target | 1.0279 | 0.0774 | |
| homer1a | Post-synaptic density (PSD) scaffolding proteins | ASD marker | 0.9026 | 0.1104 | |
| egr1 | early growth response 1 | FMRP CLIP target, ASD marker | 0.9766 | 0.0638 | |
| shank3a | SH3 and multiple ankyrin repeat domains 3 | FMRP CLIP target, ASD marker | 0.8434 | 0.1188 | |
| nrxn1 | Neurexin 3 | FMRP CLIP target, ASD marker | 1.0746 | 0.1408 | |
| nlgn3 | Neuroligin 3 | FMRP CLIP target, ASD marker | 0.8223 | 0.1693 | |
| dlgap1a | Discs large associated protein 1, scaffold proteins in the postsynaptic density | FMRP CLIP target (Increased polyA in FMRP KO) | 1.1400 | 0.1796 | |
| dlgap 1b | | FMRP CLIP target (Increased polyA in FMRP KO) | 0.8574 | 0.0253 | |
| camta 1a | Calmodulin binding transcription activator 1 | FMRP CLIP target (Increased polyA in FMRP KO) | 0.5965 | 0.0622 | * |
| surf1 | Cytochrome C Oxidase Assembly Factor | Decreased polyA in FMRP KO | 0.9191 | 0.0200 | |
| htt | Huntingtin | FMRP CLIP target | 0.6333 | 0.0758 | * |
| opa1 | Mitochondrial Dynamin Like GTPase | FMRP-Htt target | 1.2306 | 0.0958 | |
| mfn1 | Transmembrane GTPase Mitofusin 1 | FMRP-Htt target | 0.6403 | 0.0398 | ** |
| mfn2 | Transmembrane GTPase Mitofusin 2 | FMRP-Htt target | 0.6421 | 0.0103 | ** |
| drp1 | Dynamin 1 Like GTPase | FMRP-Htt target | 0.4145 | 0.2133 | |
| tfam | Transcription Factor A, Mitochondrial | Mitochondrial biogenesis | 1.1661 | 0.1665 | |
| bdnf | Brain-derived neurotrophic factor | FMRP target and regulator | 1.0448 | 0.0712 | |
| GFAP | Glial fibrillary acidic protein | Astrocyte marker | 2.6610 | 0.1676 | *** |
| olig2 | Oligodendrocyte transcription factor (OLIG2) | Oligodendrocyte marker | 1.0127 | 0.0141 | |
| aqp4 | Aquaporin4, water channel | Altered in autism | 0.4600 | 0.0400 | |

### Supplementary Figure 1

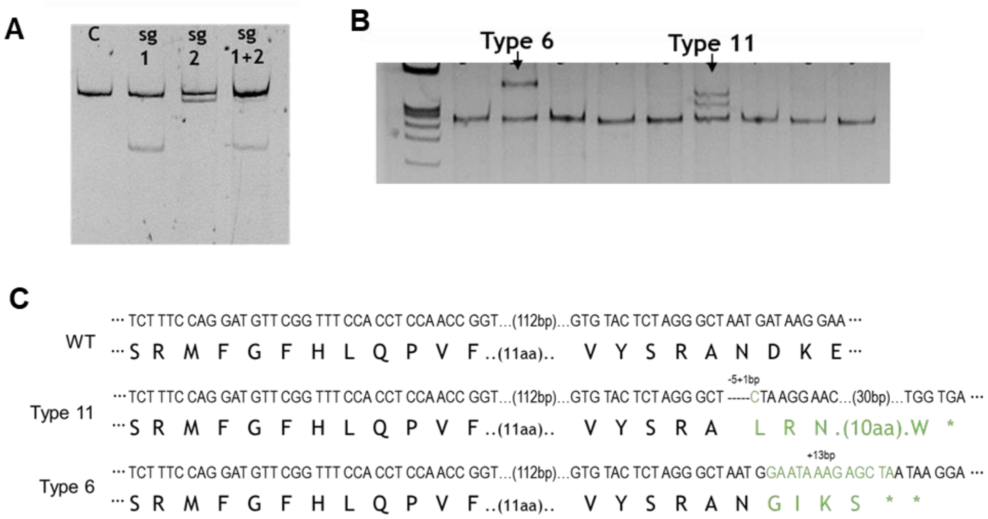


### Supplementary Figure 2

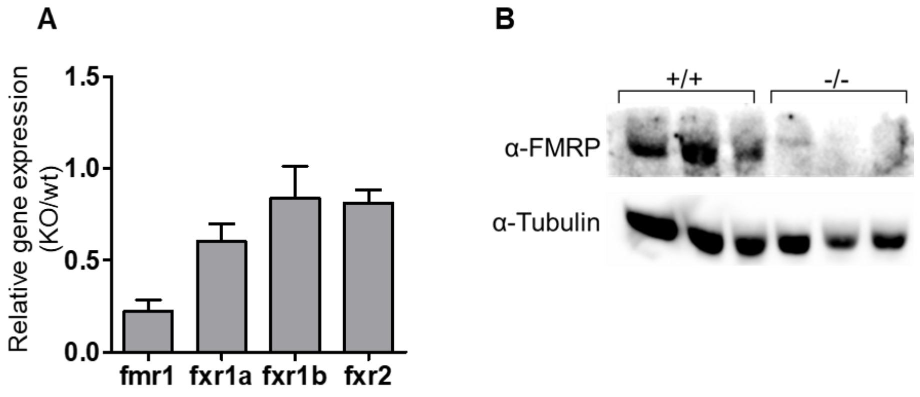

**Supplementary Figure 3**

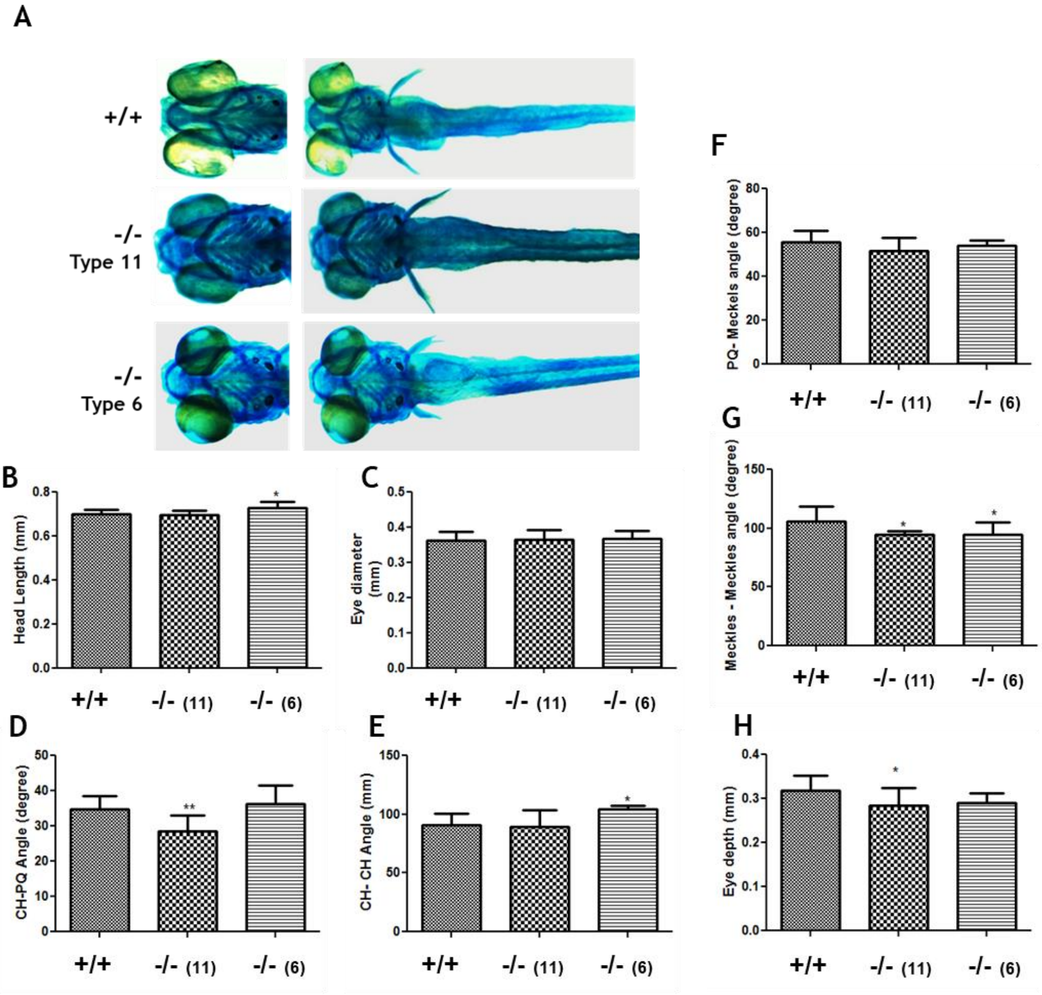

**Supplementary Figure 4**

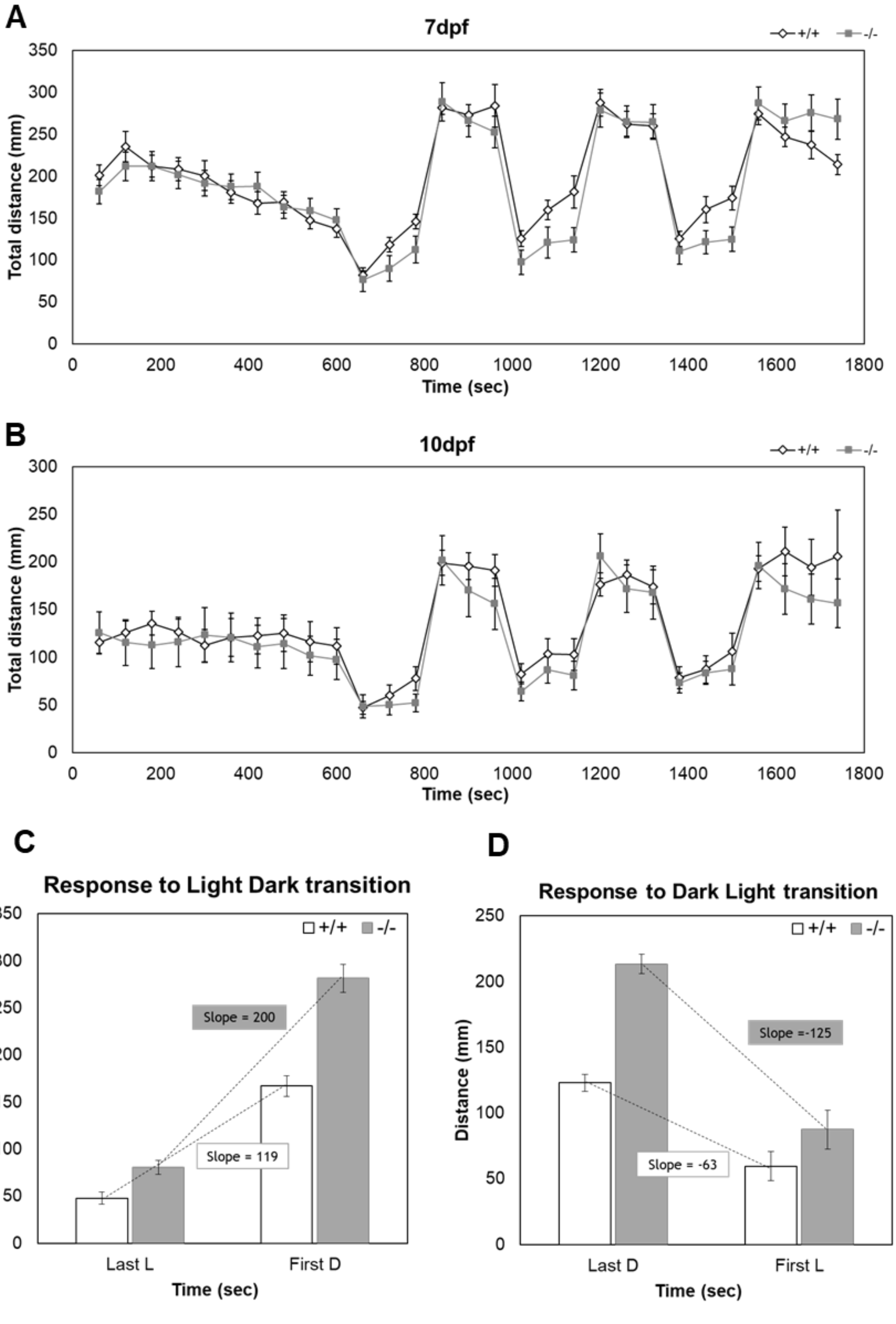

**Supplementary Figure 5**

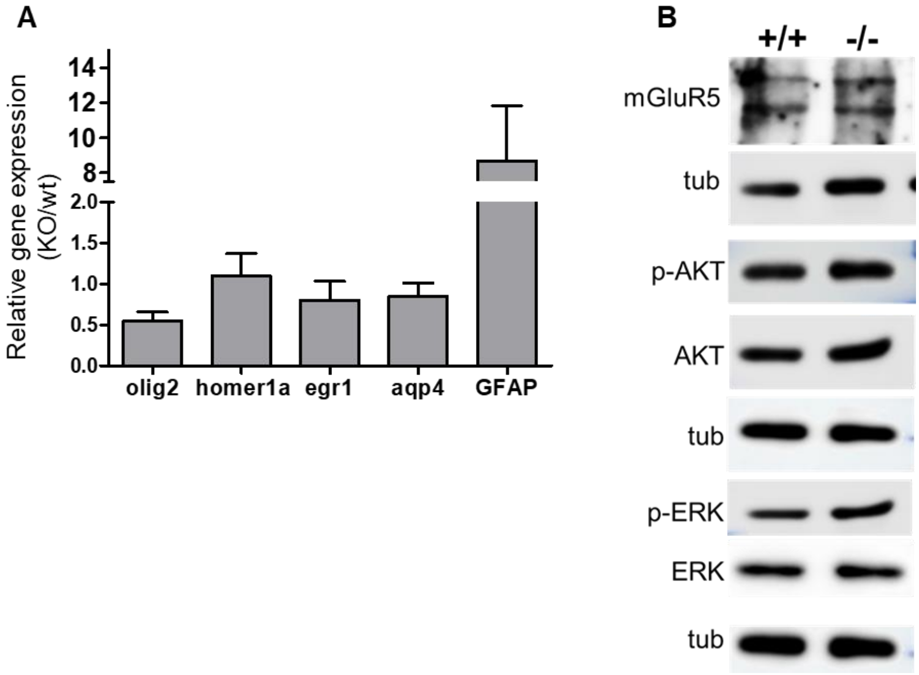


**Supplementary Figure 6**

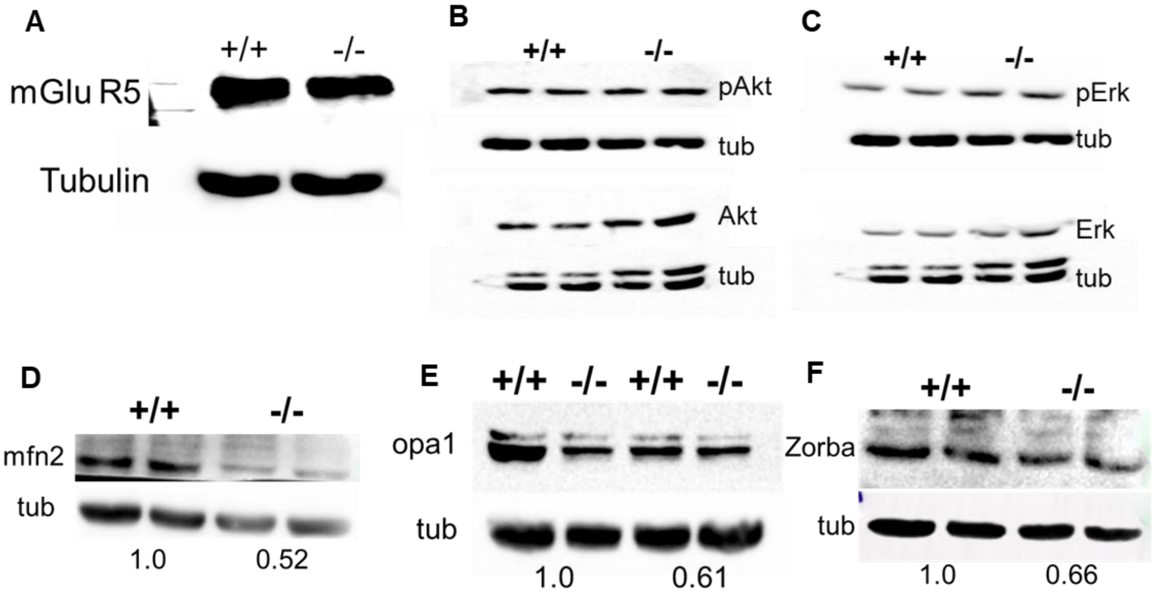

### Supplementary Figure 7

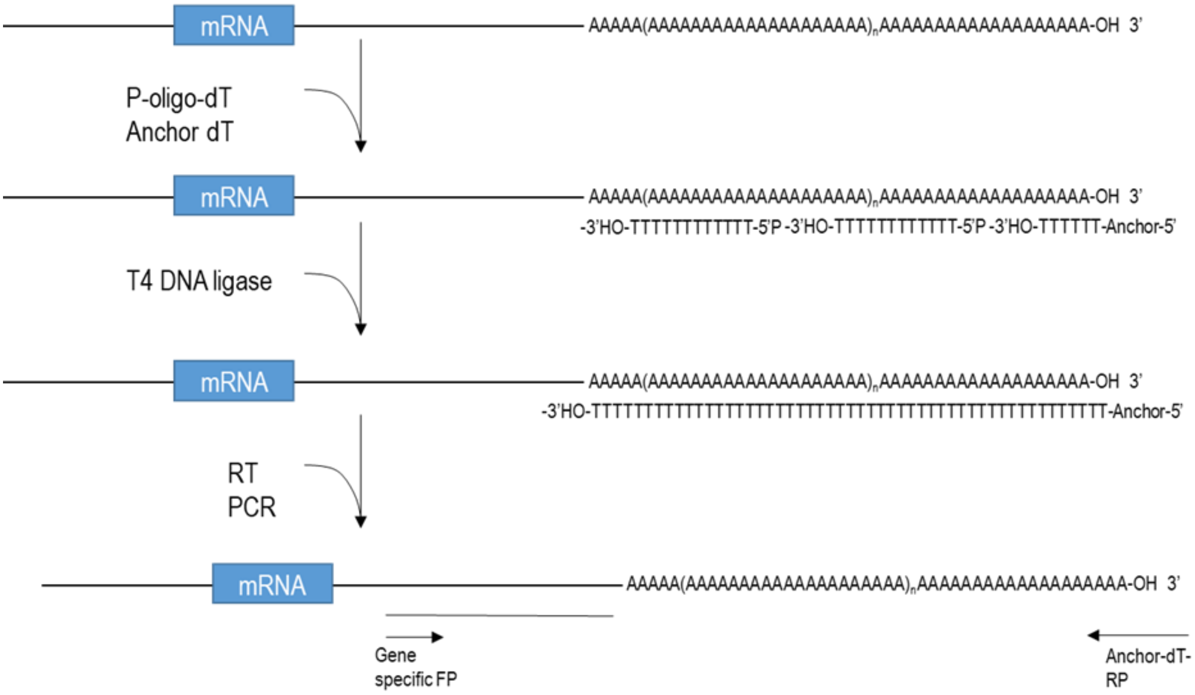


### Supplementary Figure 8

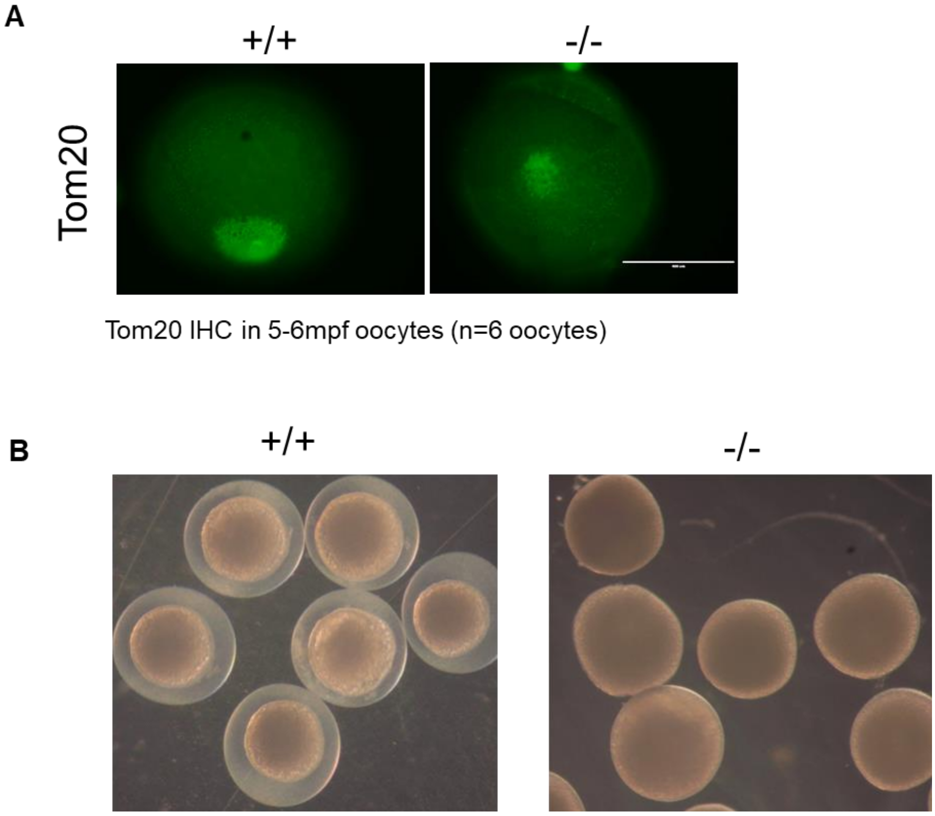

**Supplementary Figure 9**

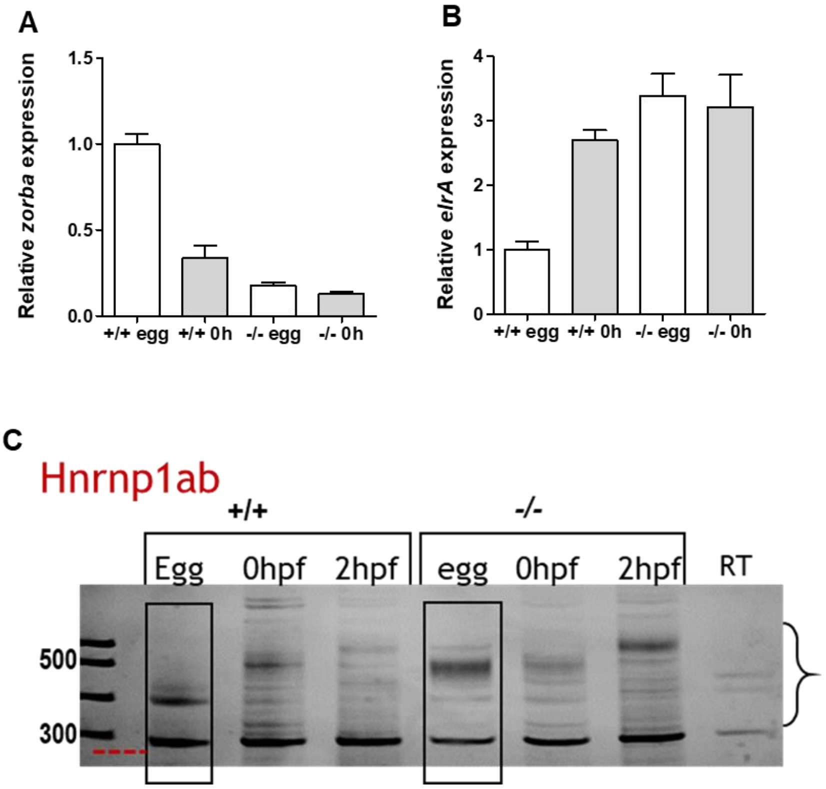

88 **Supplementary Figure 10**

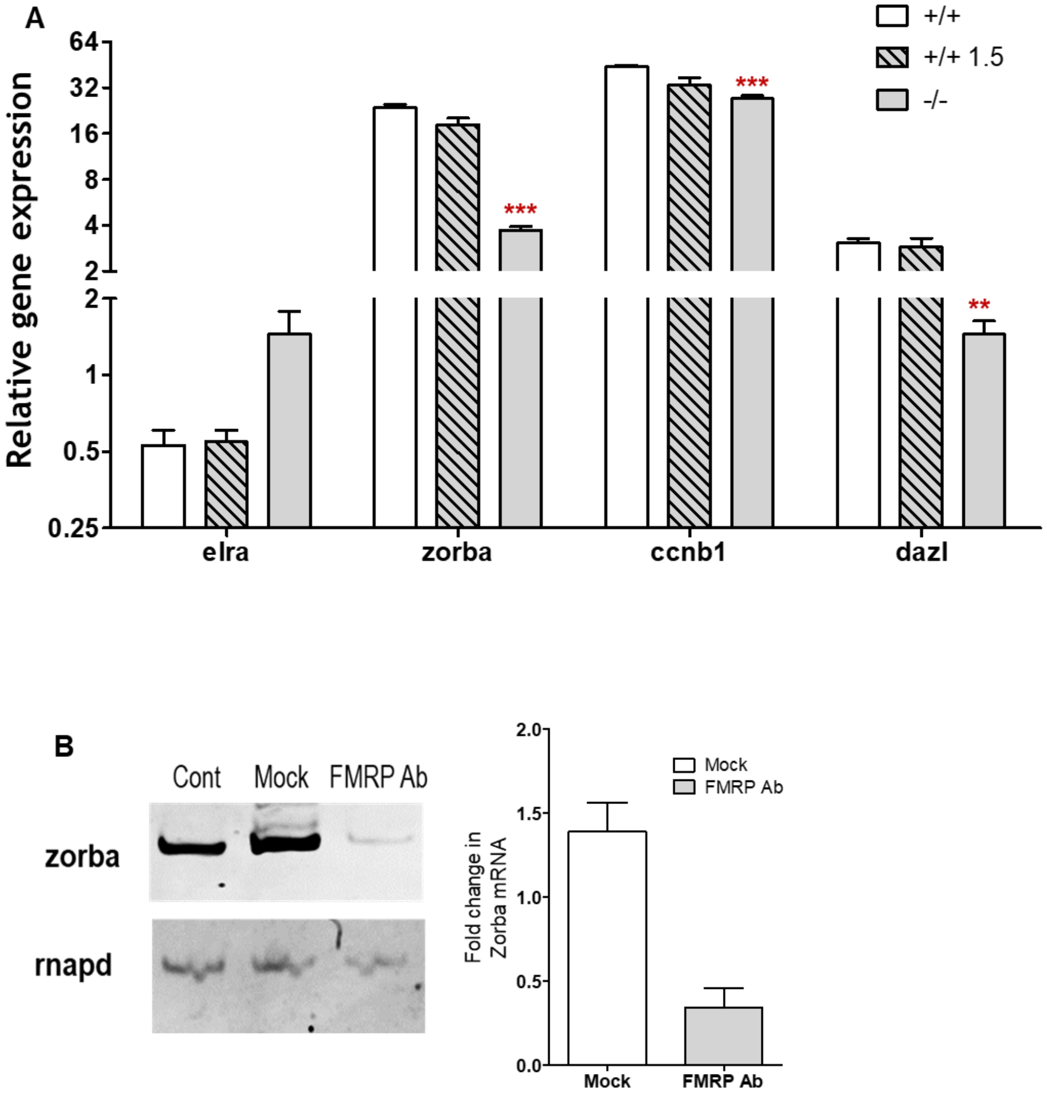


89

**Supplementary Figure 11**

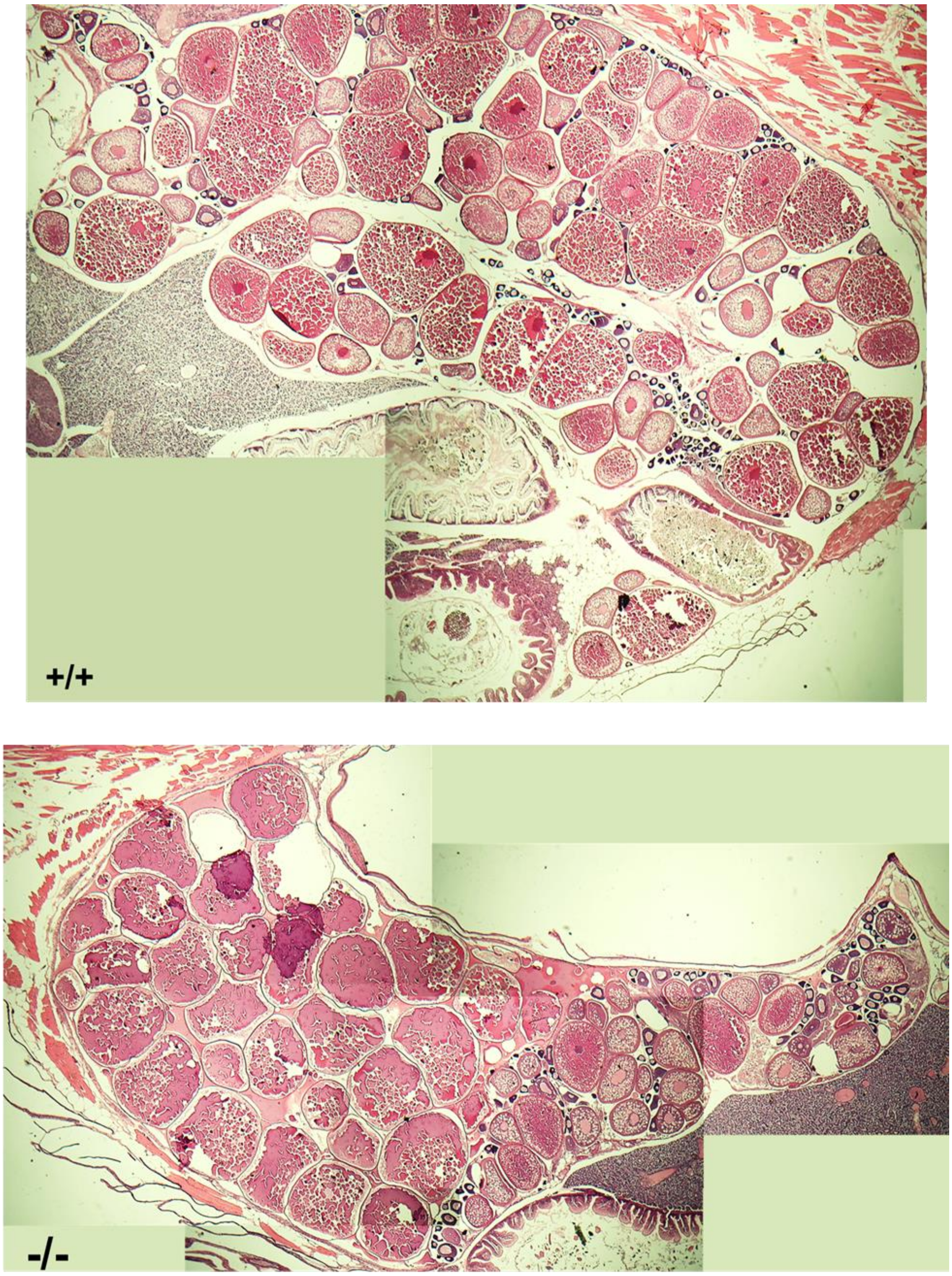
